# Adjuvant and immunomodulatory potential of *in vivo* Natural Killer T (NKT) activation by NKTT320

**DOI:** 10.1101/2021.04.27.441000

**Authors:** Nell G. Bond, Marissa Fahlberg, Shan Yu, Namita Rout, Dollnovan Tran, Taylor Fitzpatrick-Schmidt, Lesli Sprehe, Elizabeth Scheef, Joseph C. Mudd, Robert Schaub, Amitinder Kaur

## Abstract

Invariant natural killer T-lymphocytes (iNKT) are unique immunomodulatory innate T-cells with an invariant TCRα recognizing glycolipids presented on MHC class-I-like CD1d molecules. Activated iNKT rapidly secrete pro-and anti-inflammatory cytokines, potentiate innate and adaptive immunity, and modulate inflammation. Here, we report the effects of *in vivo* iNKT activation by a novel humanized monoclonal antibody, NKTT320, that binds to the invariant region of the iNKT TCR. NKTT320 led to rapid iNKT activation, increased polyfunctionality, and elevation of multiple plasma analytes within 24 hours of administration. Flow cytometry and RNA-Seq confirmed downstream activation of multiple immune subsets, enrichment of JAK/STAT and PI3K/AKT pathway genes, and upregulation of inflammation-modulating genes CMKLR1, ARG2 and NLRP12. NKTT320 also induced iNKT trafficking to adipose tissue and did not cause iNKT anergy. Our data indicate that NKTT320 has a sustained effect on *in vivo* iNKT activation, potentiation of innate and adaptive immunity, and resolution of inflammation, which supports its future use as an immunotherapeutic and vaccine adjuvant.

**Summary:** iNKTs are known immunomodulatory cells whose activation is a potential target for immunotherapies and use as an adjuvant. Here we report the potential utility of *in vivo* iNKT activation using the novel humanized monoclonal antibody NKTT320 for this purpose.

## Introduction

Invariant Natural Killer T (iNKT) lymphocytes are rare, innate T-lymphocytes with unique antigen recognition and immunomodulatory properties that make up approximately 0.01-0.1% of circulating T-lymphocytes in humans (Berzins et al., 2011). iNKT were first discovered in mice during anti-tumor studies and named for the expression of the natural killer (NK) marker NK1.1 on T-lymphocytes. iNKT differ from classical T-lymphocytes in multiple distinct, important ways. iNKT express a conserved T-cell receptor (TCR) Vα chain with an invariant complementary determining region 3 (CDR3α) as opposed to the polymorphic TCR on classical T-lymphocytes. Primate iNKT express Vα24-Jα18 which preferentially pairs with a restricted repertoire of Vβ subunits, generally Vβ11 (Bendelac et al., 2007). In contrast to classical T-lymphocytes, iNKT recognize and are rapidly activated by both endogenous and exogenous glycolipid antigens presented by antigen presenting cells (APCs) on nonpolymorphic MHC-class-I-like CD1d molecules. The classical, most widely studied iNKT activating lipid antigen is alpha-galactosylceramide (αGC), derived from the marine sponge *Agelas mauritianus* and first identified in murine cancer studies (Berzins et al., 2011). Upon activation, iNKT rapidly produce a wide range of cytokines covering T-helper (Th) 1, Th2 and Th17 functionality, often from the same cell (Van Kaer et al., 2015). iNKT in primates are more strictly defined by co-staining of the αGC-loaded CD1d tetramer (CD1dTM) and Vα24 on CD3^+^ T-lymphocytes (Watarai et al., 2008, Rout et al., 2010).

Due to their rapid response and broad functional potential, iNKT bridge the gap between innate and adaptive immunity (Brennan et al., 2013). Once activated, iNKT can be directly cytolytic (through perforin and granzyme B) and display Th1, Th2 and Th17 effector functions. Additionally, iNKT rapidly influence the function of multiple immune subsets (Brennan et al., 2013). Bidirectional interactions between iNKT and dendritic cells (DC) enhances DC maturation and facilitates antigen cross-presentation and priming of antigen-specific T-lymphocyte responses (Fujii et al., 2004, Stober et al., 2003). Activated iNKT can potentiate macrophage phagocytic function and affect polarization (Brennan et al., 2013). IFNγ production by iNKT rapidly activates natural killer (NK) cells improving cytolysis (Carnaud et al., 1999). Finally, iNKT are known to recruit and provide help to B-cells, improving B-cell maturation, antibody class-switching and overall humoral immunity (Chang et al., 2011, King et al., 2011). Due to their diverse immunomodulatory properties, there is great interest in harnessing iNKT activation as an immunotherapeutic tool and a vaccine adjuvant.

Studies exploring αGC-mediated iNKT activation as a vaccine adjuvant have largely been conducted in mice with varying degrees of success (Kopecky-Bromberg et al., 2009, Gonzalez-Aseguinolaza et al., 2002, Fujii et al., 2003, Silk et al., 2004, Venkataswamy et al., 2009). One barrier to the use of iNKT activating agents such as soluble αGC *in vivo* is subsequent iNKT anergy in which iNKT are rendered unable to respond to further stimuli (Fujii et al., 2002, Parekh et al., 2005). Although administration of αGC loaded on autologous DCs has shown promise for cancer immunotherapy in human studies (Kunii et al., 2009, Motohashi et al., 2011, Chang et al., 2005), there is a need for alternate strategies of *in vivo* iNKT activation that can effectively harness the immunomodulatory properties of iNKT for widespread therapeutic use.

Antibodies directed against the iNKT cell receptor are one such class of potential alternative iNKT modulating agents. NKTT120 is a humanized monoclonal iNKT depleting antibody developed by NKT Therapeutics (Sharon, MA) that directly binds to the CDR3 region of the Vα−subunit of the semi-invariant iNKT TCR with high affinity (Scheuplein et al., 2013). NKTT120 was engineered with an IgG1 Fc, thus supporting Fc-receptor binding and iNKT depletion by antibody dependent cellular cytotoxicity (ADCC) (Scheuplein et al., 2013). NKTT120 successfully depleted iNKT without any adverse effects in healthy humans and macaques (Scheuplein et al., 2013, Field et al., 2017). The humanized monoclonal antibody NKTT320 developed by NKT Therapeutics shares the variable region and iNKT binding specificity with NKTT120 but was engineered with an IgG4 Fc (Scheuplein et al., 2013) allowing it to successfully bind and activate iNKT without ADCC-mediated depletion (Reddy et al., 2000, Patel et al., 2020). *In vitro* iNKT activation in response to NKTT320 has been characterized previously (Patel et al., 2020), however, the *in vivo*, translational potential has yet to be described.

In this study, we characterized the *in vivo* effects of NKTT320 administration on iNKT function and bystander lymphocyte subsets in Mauritian-origin cynomolgus macaques (MCM). We show rapid induction of iNKT activation and polyfunctionality without anergy, downstream effects on monocytes, T- and B-lymphocytes, and gene enrichment in the inflammatory response, heme metabolism, JAK/STAT signaling and PI3K/AKT pathways. Our results demonstrate the utility of NKTT320 as an *in vivo* immunomodulatory tool and novel vaccine adjuvant with high translational promise.

## Results

### NKTT320 specifically activates iNKT

To characterize the potential of NKTT320 as an adjuvant, we first conducted *in vitro* studies in unfractionated peripheral blood mononuclear cells (PBMCs) of MCM. MCM were chosen because their circulating iNKT frequencies and phenotype are similar to humans (Rout et al., 2012). As previously reported, iNKT were identified by flow cytometry as T-lymphocytes co-expressing the Vα24 TCR and binding to αGC-loaded CD1d-tetramers (CD1dTM) (Rout et al., 2010, Rout et al., 2012) (Figure 1A, Supplemental Figure 1A). PBMCs stimulated *in vitro* with goat-anti-mouse-IgG (GAM-IgG) cross-linked NKTT320 at 200ng/mL showed specific iNKT activation as evidenced by CD3 downregulation, CD69 upregulation, and secretion of IFNγ and TNFα compared to media alone (Figure 1B, Supplemental Figure 2A). Escalating concentrations of NKTT320 showed a dose-dependent effect on iNKT activation as indicated by downregulation of Vα24, CD1dTM, and CD3 (Figure 1C-D, Supplemental Figure 2B). At higher doses, a combination of TCR downregulation and competition for the same invariant TCR binding site resulted in loss of sensitivity to detect CD1dTM-positive iNKT (Figure 1C). iNKT activation was not associated with non-NKT T-lymphocyte activation during a 24-hour stimulation period demonstrating the iNKT specificity of NKTT320 (Figure 1B, D).

**Figure 1.**
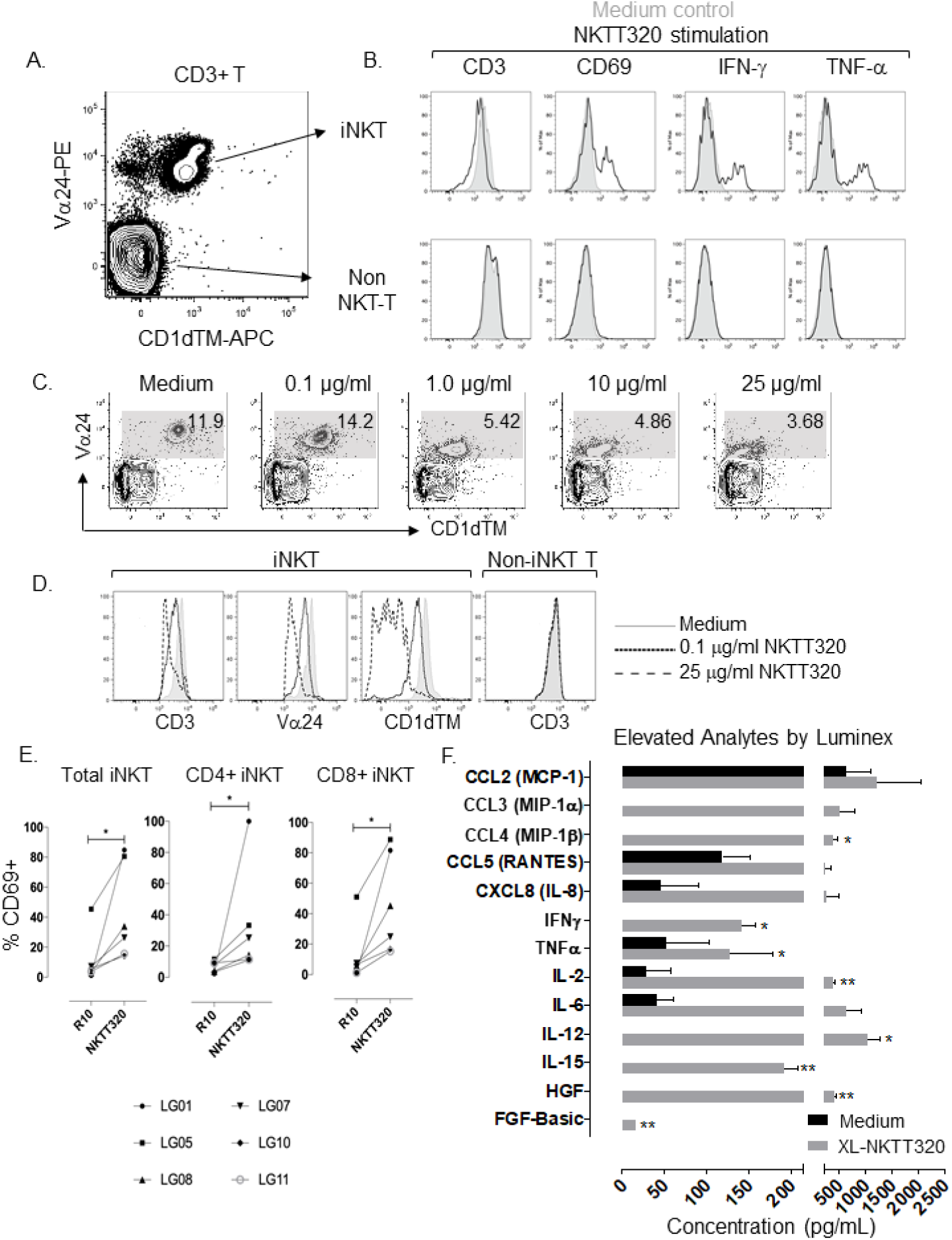
iNKTs are specifically activated by NKTT320. (A) Representative plot showing iNKT and non-NKT T-cell gating strategy gated on CD3+ T-cells. (B) Overlay histograms (unstimulated=shaded, NKTT320=open histogram) comparing CD3, CD69 and cytokine expression on R10 and NKTT320 (200ng/mL) treated PBMCs stimulated for 4hrs *in vitro*, iNKTs (top) and non-NKT T–cells (bottom). (C-D) *In vitro* data showing iNKT frequency and TCR downregulation after overnight incubation with escalating concentrations of NKTT320. TCR downregulation is measured by Vα24, CD1d and CD3. (E) CD4+ and CD8+ iNKT activation *in vitro* measured by CD69 in 6 animals stimulated with 200ng/mL NKTT320 for 4 hours. (F) Luminex data showing analytes which were elevated in cells stimulated with 200ng/mL NKTT320 compared to medium. Supernatants were collected after 48 hours. Statistics were done by non-parametric Wilcoxon signed-rank test (n=3). *<0.05, **<0.01.

We then evaluated *in vitro* NKTT320 treatment for differences in activation of single positive CD4 or CD8 iNKT. Unstimulated cultured CD4^+^ and CD8^+^ iNKT did not differ in baseline activation levels measured by surface CD69 and HLA-DR expression (Figure 1E and data not shown). NKTT320 treatment led to significant activation of both CD4^+^ (p=0.0312) and CD8^+^ iNKT (p=0.0312) as measured by CD69 upregulation (Figure 1E). There was no significant difference in activation of CD4^+^ compared to CD8^+^ iNKT (data not shown).

To examine broader functional changes resulting from NKTT320 treatment we cultured PBMC *in vitro* for 48 hours with either media alone (mock stimulation) or with cross-linked NKTT320 and measured secreted analytes using a non-human primate (NHP) 29-plex Luminex (Figure 1F). Eight analytes significantly elevated in the NKTT320-treated supernatants included proinflammatory and Th1 cytokines (IFNγ, TNFα, IL-12, IL-2, IL-15), chemokines (CCL4), and growth factors (FGF-basic, HGF). IL-6, CXCL8 (IL-8), CCL2, CCL3, and CCL5 (RANTES) trended higher but the increase did not reach statistical significance.

### Differential iNKT activation with NKTT320 and αGC stimulation

Since glycolipid agonists are being used for iNKT activation, we compared NKTT320 to the classic iNKT agonist, αGC. In a series of time course *in vitro* stimulation experiments, all parameters of iNKT activation (CD69, IFNγ, TNFα, IL-2 and IL-4) appeared more rapidly and were of greater magnitude after NKTT320 as compared to αGC stimulation (Figure 2A). Increased IFNγ and TNFα secretion, and increased iNKT polyfunctionality were apparent within 4 hours of NKTT320 stimulation (Figure 2A-B). With the exception of IL-4, αGC-stimulated iNKT were able to match responses from NKTT320-stimulated cells by 48 hours (Figure 2A). The kinetic and qualitative differences in iNKT response between NKTT320 and αGC stimulation has implications for therapeutic choice of NKT agonist.

**Figure 2.**
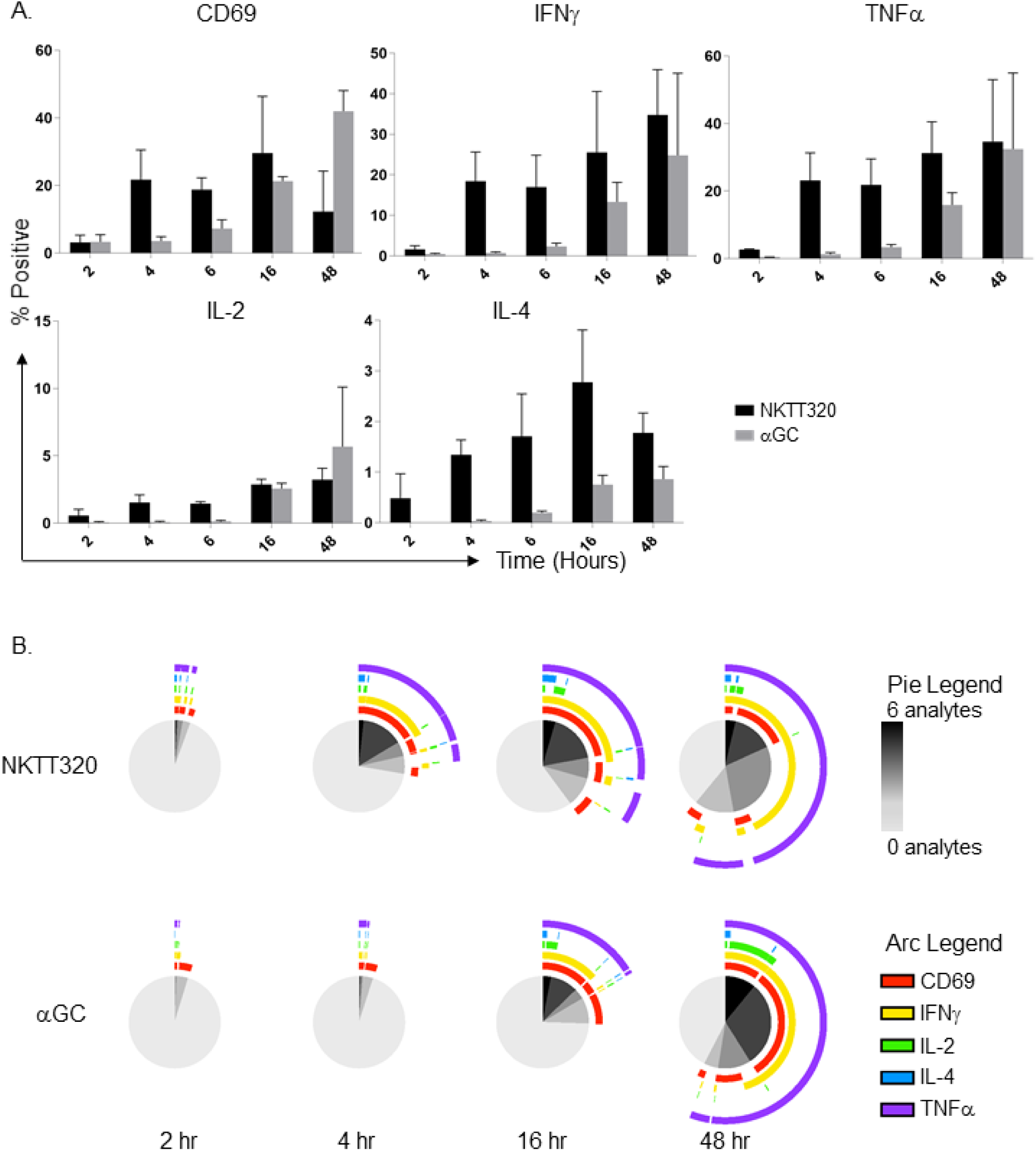
Differential kinetics of NKTT320 and αGC activation *in vitro*. (A) Intracellular cytokine staining of PBMCs stimulated with goat-anti-mouse-IgG cross-linked NKTT320 or αGC loaded on C1Rd cells, harvested and stained at 2, 4, 6, 16, and 48 hours. Data on cytokine secretion gated on Vα24+ cells. Mean and SEM shown (n=2). (B) SPICE analysis of experiment described in (A) showing kinetic differences in speed, magnitude and co-expression of cytokines in NKTT320 and αGC stimulated iNKT cells (n=2).

### NKTT320 Pharmacokinetics

After determining the effectiveness and specificity of iNKT activation with NKTT320, we set out to determine the pharmacokinetics of this antibody following *in vivo* administration. Dose escalation experiments were performed in three groups of MCM administered a single IV dose of NKTT320 at low (100μg/kg, n=5), mid (300μg/kg, n=3) and high (1000μg/kg, n=3) concentrations. These doses were selected based on previously published data of NKTT120 (Scheuplein et al., 2013, Field et al., 2017). No adverse effects were observed following *in vivo* NKTT320 administration. Peak serum NKTT320 levels ranging between 5.94-77.46μg/mL were reached within 24 hours of a single IV dose. The iNKT TCR saturation level previously determined for NKTT120 on a Biacore assay against the immobilized iNKT TCR resulted in a K_D_ of 44nm corresponding to a saturation level of 6μg/mL (Scheuplein et al., 2013). All NKTT320-treated animals in the high-and mid-dosing groups reached peak levels above the NKT TCR saturation level (6μg/mL) within 30 minutes of antibody administration while 4 of five animals in the low dose (100μg/kg) surpassed TCR saturation levels within 2 hours of administration (Figure 3A).

**Figure 3.**
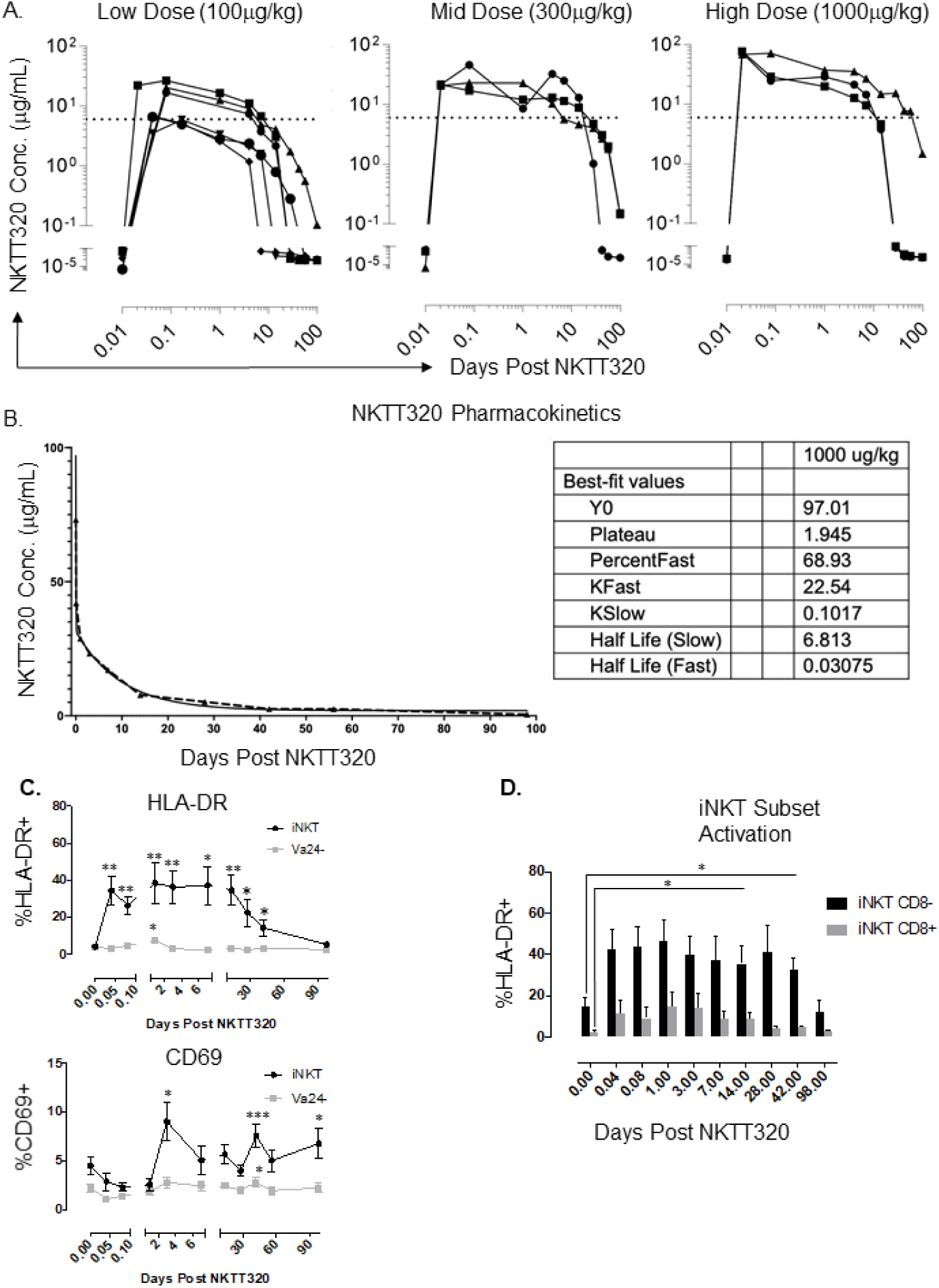
*In vivo* iNKT activation and pharmacokinetics of NKTT320. (A) Plasma NKTT320 concentration measured by ELISA in animals that received Low (100ug/kg, n=5), Mid (300ug/kg, n=3) and High (1000ug/kg, n=3) doses of NKTT320 IV. Horizontal dotted line shows NKT TCR saturation level (6ug/mL). Samples were taken at day 0 and 30 minutes, 2 hours, day 1, 4, 7, 14, 28, 42, 56, and 98 post-NKTT320 treatment. (B) NKTT320 pharmacokinetic data determined by a single dose of 1000ug/kg. Half-life shown in days. (C) Mean HLA-DR and CD69 expression in iNKT versus non-iNKT-T. (D) iNKT CD8- vs iNKT CD8+ in animals treated *in vivo* with NKTT320 (n=9). Paired t-tests were used to determine statistical significance of increased activation of total iNKT or Va24-T (C), and CD8+ or CD8-iNKT (D) compared to their respective baseline values. * <0.05.

The kinetics of circulating NKTT320 showed animal to animal variation. Peak NKTT320 serum levels trended higher in the high dose group but did not reach significance (Figure 3A and data not shown). Plasma NKTT320 levels declined below detection in all animals 2-6 weeks post treatment, excepting one animal in the high dose group that maintained detectable serum NKTT320 levels through week 14. NKTT320 plasma concentrations in the high dose group were used to determine the half-life of NKTT320 (Figure 3B). The calculated half-life of 6.81 days or 163.2 hours post administration is less than the half-life of NKTT120 but consistent with variation in plasma clearance of human IgG antibodies in NHPs (Scheuplein et al., 2013, Newman et al., 2001).

One multiply dosed animal with successive NKTT320 doses administered at two-week intervals (dose 1=30μg/kg, dose 2=100μg/kg and dose 3=100μg/kg) reached the 6μg/mL NKT TCR saturation level only after the second and third dose indicating that the first dose was not sufficient to surpass TCR saturation (data not shown). Multiple doses were additive in this animal and again peaked within 24 hours after each administration (data not shown). These limited data suggest that doses above 30μg/kg NKTT320 are needed for TCR saturation.

### NKTT320 effect on iNKT after *in vivo* administration

We monitored *in vivo* changes in iNKT and non-iNKT (Vα24^−^) T-lymphocytes from 30 minutes onwards following a single intravenous dose of NKTT320 (Figure 3C-D, Supplemental Figure 3). Similar to *in vitro* stimulation (Figure 1A-D), *in vivo* NKTT320 resulted in downregulation of Vα24 TCR and CD1dTM-positive iNKT-cells within 24 hours of administration (Supplemental Figure 3A). Because NKTT320 binds to the invariant region of the NKT TCR, the decrease in CD1dTM-positive iNKT could represent a ‘masking’ effect. However, the attendant downregulation of Vα24 TCR and CD3 (data not shown) indicates that the decline in detectable Vα24/CD1dTM co-expressing iNKT following NKTT320 administration was also a result of *in vivo* NKT activation. Due to the loss of visualization of Vα24^+^CD1dTM^+^ iNKT we also enumerated total Vα24^+^ T-lymphocytes to monitor changes in circulating iNKT frequency post NKTT320 administration (Supplemental Figure 3B).

Both Vα24/CD1dTM co-staining iNKT and total Vα24^+^ T-lymphocyte frequencies declined significantly after NKTT320 administration (Supplemental Figure 3B). While Vα24/CD1dTM co-staining iNKT frequency was reduced by day 1 and remained significantly lower than baseline through week 14 (p<0.05 at all sampled time-points), a significant decline in total Vα24^+^ T-lymphocyte frequency was only observed for one week post NKTT320 administration. The decline in Vα24^+^CD1dTM^+^ iNKT, likely the result of a masking effect, persisted well beyond the time that serum NKTT320 levels were undetectable. This may mean that iNKT-bound NKTT320 undetectable in the blood is slowly released in the tissues and continues to activate iNKT.

Monitoring of HLA-DR and CD69 surface expression revealed rapid, sustained iNKT-specific activation without general T-lymphocyte activation following NKTT320 administration (Figure 3C). By 30 minutes, iNKT were significantly activated above baseline levels (p=0.0156) while non-iNKT (Vα24^−^) T-lymphocytes were not activated. iNKT then remained significantly activated for 6 weeks after treatment. Non-iNKT T-lymphocytes showed a transient significant increase in HLA-DR or CD69 expression post NKTT320 treatment. Activation levels of circulating iNKT remained significantly above non-iNKT T-lymphocytes starting within 30 minutes of NKTT320 administration (p=0.0039) and persisted for at least 14 weeks (p=0.0391).

To distinguish responses of iNKT subsets, we investigated differences in CD8^−^ and CD8^+^ iNKT activation. Both subsets were significantly activated within 30 minutes of NKTT320 administration; while CD8^−^ iNKT remained significantly activated through week 6 post treatment, increased CD8^+^ iNKT activation lasted two weeks (Figure 3D). In contrast to *in vitro* data, circulating CD8^−^ iNKT were significantly more activated compared to CD8^+^ iNKT pre- and post NKTT320 administration (p=<0.01).

### NKTT320 rapidly modulates T-lymphocyte function

Central to investigation of NKTT320’s utility as an adjuvant is its effect on other immune cell subsets. We used intracellular cytokine staining (ICS) flow cytometry to evaluate NKTT320-induced functional changes in mitogen responsiveness of iNKT and non-iNKT T-lymphocytes at discrete time-points after IV NKTT320 administration. Increased responsiveness of iNKT to overnight PMA/ionomycin stimulation was observed within 30 minutes of NKTT320 administration (Figure 4A-B). A significant increase in IL-2 secretion detected at 30 minutes post NKTT320 was sustained through day 14 post treatment. Likewise, IL-4 was significantly upregulated within two hours of NKTT320 treatment. Significant increases in IFNγ and TNFα were observed day 3 onwards. Increased responsiveness to mitogen stimulation was also observed in Vα24^−^ “non-iNKT” T-lymphocytes but at a later onset suggesting downstream activation of other T-lymphocyte subsets. The increase in cytokine production in both iNKT and non-NKT T-lymphocytes mainly originated from CD4^+^ T lymphocytes (Supplemental Figure 4A and data not shown).

**Fig. 4.**
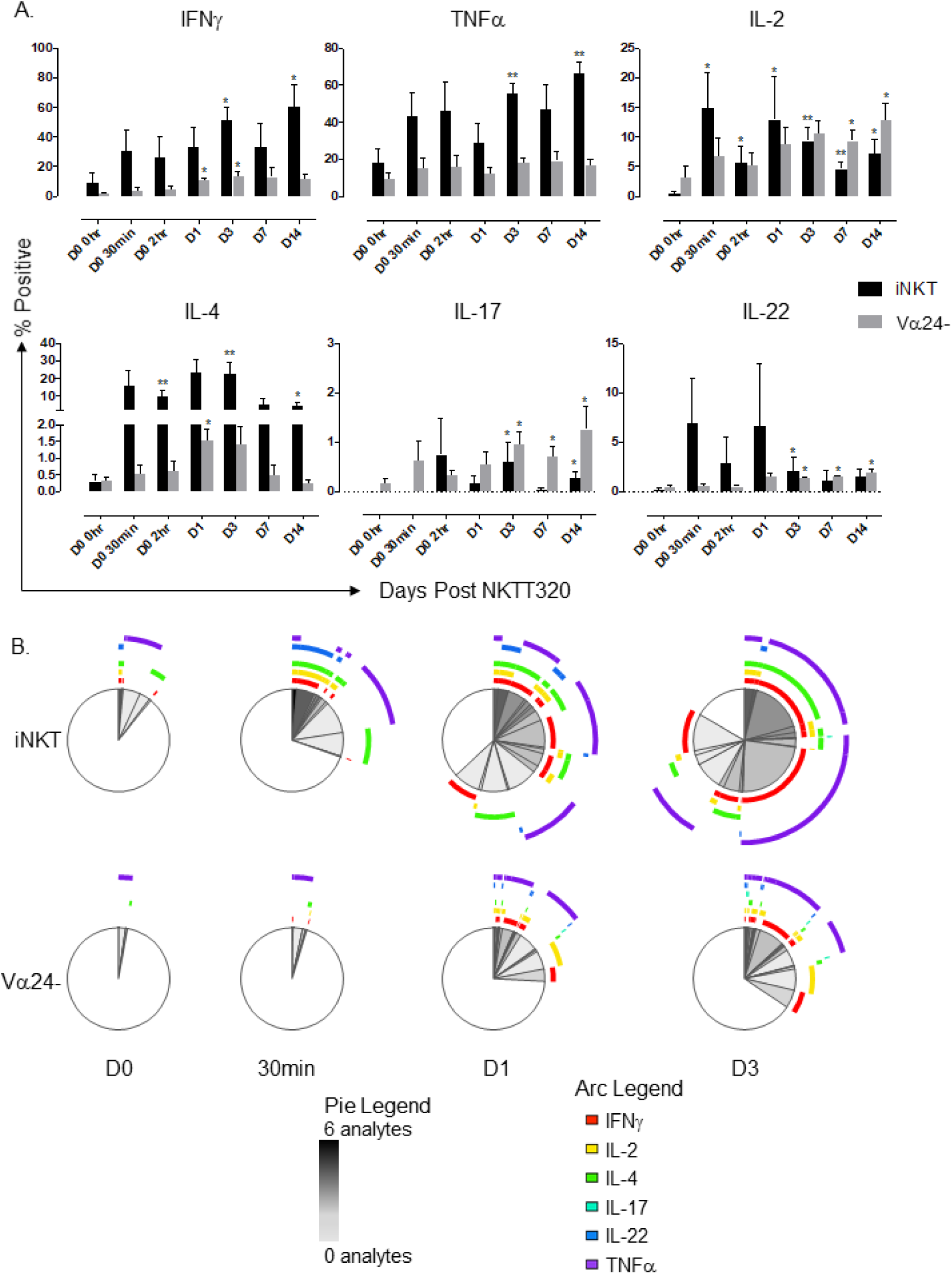
NKTT320 rapidly induces functional changes in T-cell subsets. (A) Mean cytokine expression in iNKTs and Non-iNKT Ts measured by intracellular cytokine staining following overnight PMA/ionomycin stimulation. Samples were taken at day 0 and 30 minutes, 2 hours, day 1, 4, 7, 14. Data in n=4 MCM. Significance compared to baseline is indicated by an asterisk and was determined by paired parametric t-test. *<0.05, **<0.01. (B) SPICE analysis of iNKT vs Non-iNKT T-cells indicating cytokine co-expression after stimulation with PMA/ionomycin (n=4).

NKTT320 also rapidly increased iNKT polyfunctionality; by day 1, over 50% of responding iNKT secreted two or more cytokines on mitogen stimulation (Figure 4B, Supplemental Figure 4B). Despite TCR downregulation, a discernable iNKT population was visible at early time-points post NKTT320 administration in most animals to allow evaluation of its function by flow cytometry (Supplemental Figure 4C). Increased polyfunctionality was less evident in the non-NKT T-lymphocytes. Overall, iNKT were activated rapidly in response to NKTT320 treatment and appear to have also induced changes in the functional potential of non-iNKT T-lymphocytes.

### *In vivo* iNKT activation results in rapid activation of the innate and adaptive immune system

The ability to modulate innate and adaptive immune responses is crucial when identifying promising immune modulatory tools and vaccine adjuvants. To assess the effect on the host immune response we longitudinally measured plasma analytes by Luminex, and monitored changes in immune cell frequency, phenotype and function by flow cytometry after *in vivo* administration of NKTT320.

A significant increase in 21 plasma analytes was detected in the first 72 hours of NKTT320 administration (Figure 5A, Supplemental Table 1). Plasma CCL2, CXCL10 and IL-6 were elevated within 30 minutes reaching peak levels in the first 2 hours (Figure 5B). Significantly elevated analytes that peaked in the first 24 hours included cytokines and chemokines that likely originated from iNKT (IL-2, IL-4, IL-5, IL-6, IL-10, CCL4, CCL5) and from downstream activation of dendritic cells and monocytes/macrophages (IL-12, IL-1β, IL-1RA, CXCL8, CXCL9-11, IL-6, CCL2, CCL22). Interestingly, IFNγ, TNFα and GM-CSF peaked later at 72 hours (Supplemental Table 1) suggesting iNKT and non-iNKT sources such as NK cells and other T-lymphocytes. Of note, the increase in plasma analytes corresponded temporally to the peak of plasma NKTT320 and iNKT-specific activation. The early response was indicative of rapid activation of the innate immune system and included proinflammatory analytes as well as several chemo-attractants crucial for immune cell recruitment of monocytes, granulocytes, NK cells and activated T-lymphocytes (Bromley et al., 2008, Deshmane et al., 2009).

**Fig. 5.**
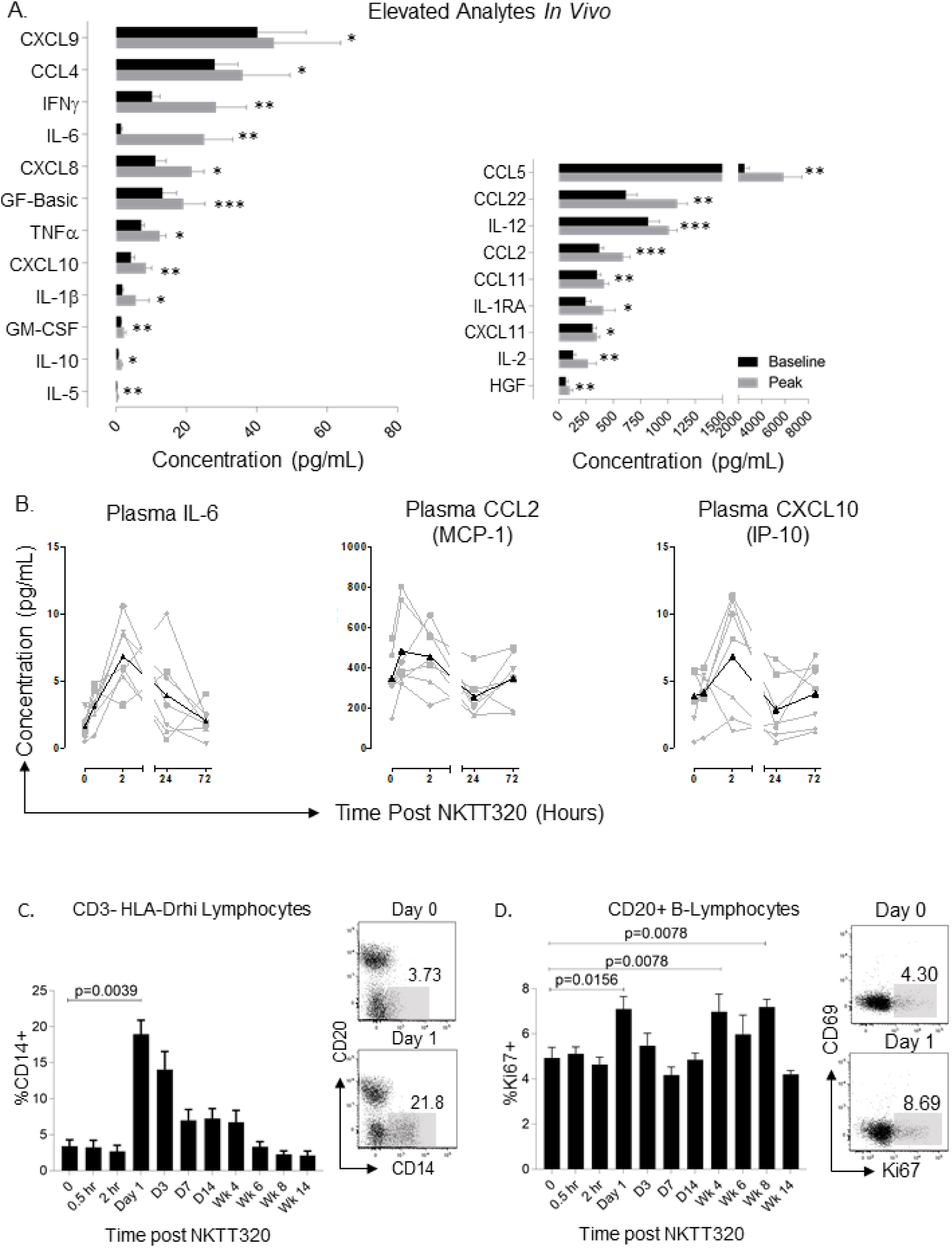
NKTT320 rapidly induces functional changes *in vivo*. (A) Analytes significantly upregulated after NKTT320 treatment *in vivo* measured by plasma Luminex at baseline and peak (n=12). *<0.05, **<0.01, ***<0.001.(B) *Ex vivo* plasma Luminex kinetics post NKTT320 for IL-6, CCL2, and CXCL10 in 7 animals from a single Luminex run. (C) Monocyte frequency shown as percentage of CD3-HLA-DRhi lymphocytes. Representative plots show an increase in circulating monocytes on day 1 compared to day 0. (D) B-cell proliferation measured by Ki67 expression on CD20+ B-cells. Representative plots show increases in B-cell Ki67 expression at day 1 post NKTT320 administration. Significant differences from baseline were determined through non-parametric Wilcoxon signed-rank test (n=9 MCM).

Consistent with the Luminex data, flow cytometry revealed significant changes in circulating monocytes and B-lymphocytes (Figure 5C-D). A significant increase in CD14^+^ monocytes detected at day 1 post NKTT320 treatment had declined to baseline levels by week 6 (Figure 5C). Increased levels of proliferating CD20^+^ B-lymphocytes as measured by Ki67 were detected beginning at day 1 (p=0.0156) and intermittently thereafter (Figure 5D). These data are consistent with downstream effects of iNKT activation on other immune cell populations. The broad range of cytokines, chemokines and growth factors detected in response to NKTT320 treatment underscores its ability to rapidly activate the innate immune system and potentially serve as a vaccine adjuvant.

### RNA-Seq analysis of effects of *in vivo* NKTT320

To further investigate host immune modulation by NKTT320, we performed bulk RNA-Seq analysis on frozen unfractionated PBMC collected at 0, 30 minute, 2-hour, and 24-hour time-points post NKTT320 administration in four MCM with iNKT frequencies ranging between 0.1-10% of circulating T-lymphocytes (Figures 6-7). RNA-Seq analysis revealed significant sequential differential gene expression following NKTT320 treatment. At 30 minutes post NKTT320, there were 104 unique genes differentially expressed, followed by 244 and 740 genes at 2 and 24-hours, respectively (Figure 6A, Supplemental Table 2). Volcano plots showed significant changes in several genes. Notably, the scavenger receptor CD163 exclusively expressed on monocytes and macrophages (Moestrup and Moller, 2004), was significantly upregulated as early as 30 minutes and sustained through 24 hours (Figure 6B and 6C). Concomitantly, CD83 and TNF-Alpha-Induced Protein 3 (TNFAIP3) were significantly downregulated at 24-hours (Figure 6B). CD83, a member of the immunoglobulin (Ig) superfamily expressed on mature DCs can regulate inflammation by suppressing IL-12 and MCP-1 production (Bates et al., 2015). Downregulation of TNFAIP3 or A20, an inhibitor of TCR signaling and NF-κB activation (Das et al., 2018) was consistent with promotion of an inflammatory response.

**Fig. 6.**
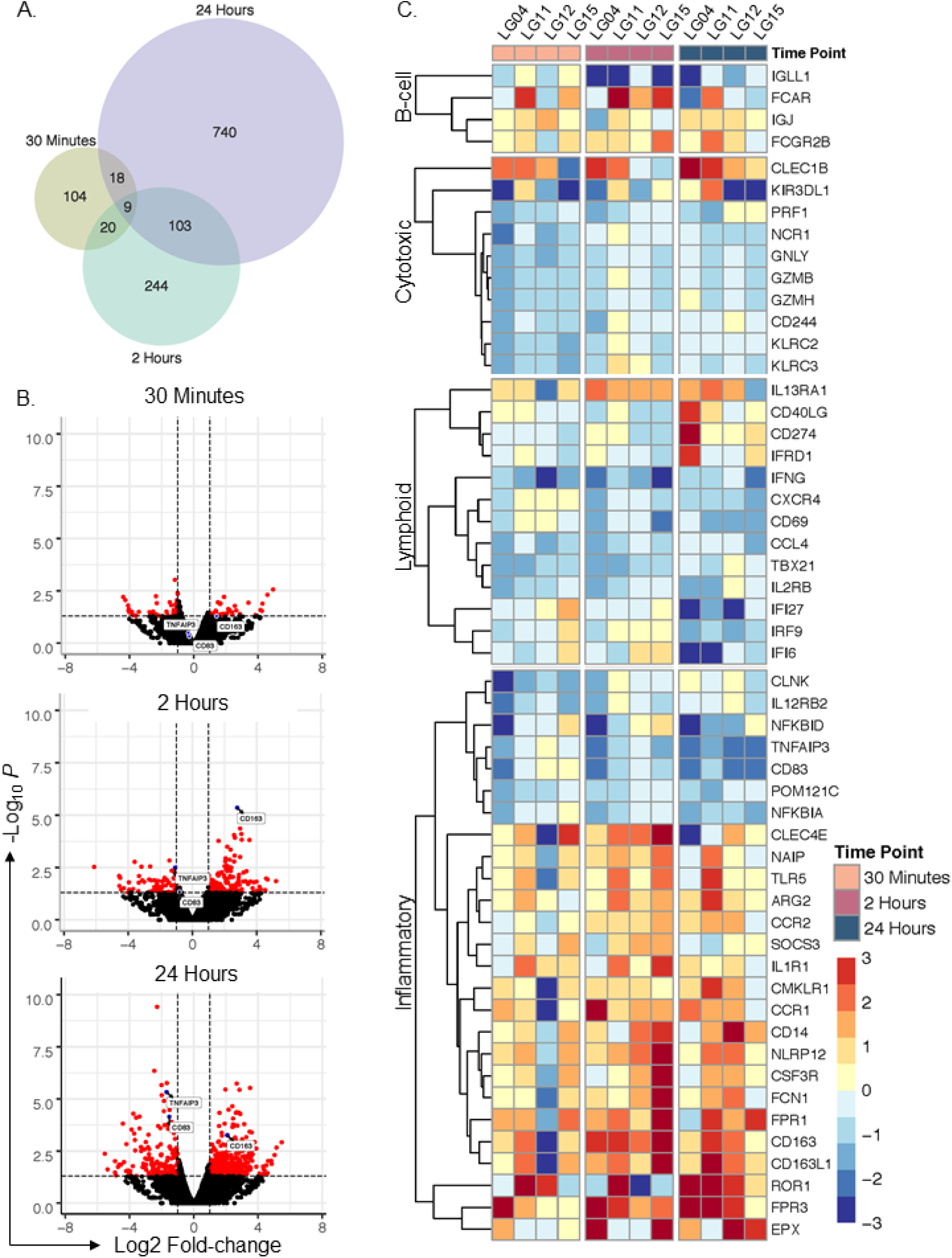
RNA-seq analysis reveals rapid differential gene expression. A) Venn diagram showing differentially expressed genes where p-value <0.05 at 30 minutes, 2 hours and 24 hours. (B) Volcano plots of all genes showing kinetic increases in transcriptomic changes over time. (C) Heatmaps showing differential expression (log_2_ fold change) for targeted genes where p-value <.05 related to B-cell, cytotoxic, lymphoid and inflammatory pathways.

**Figure 7.**
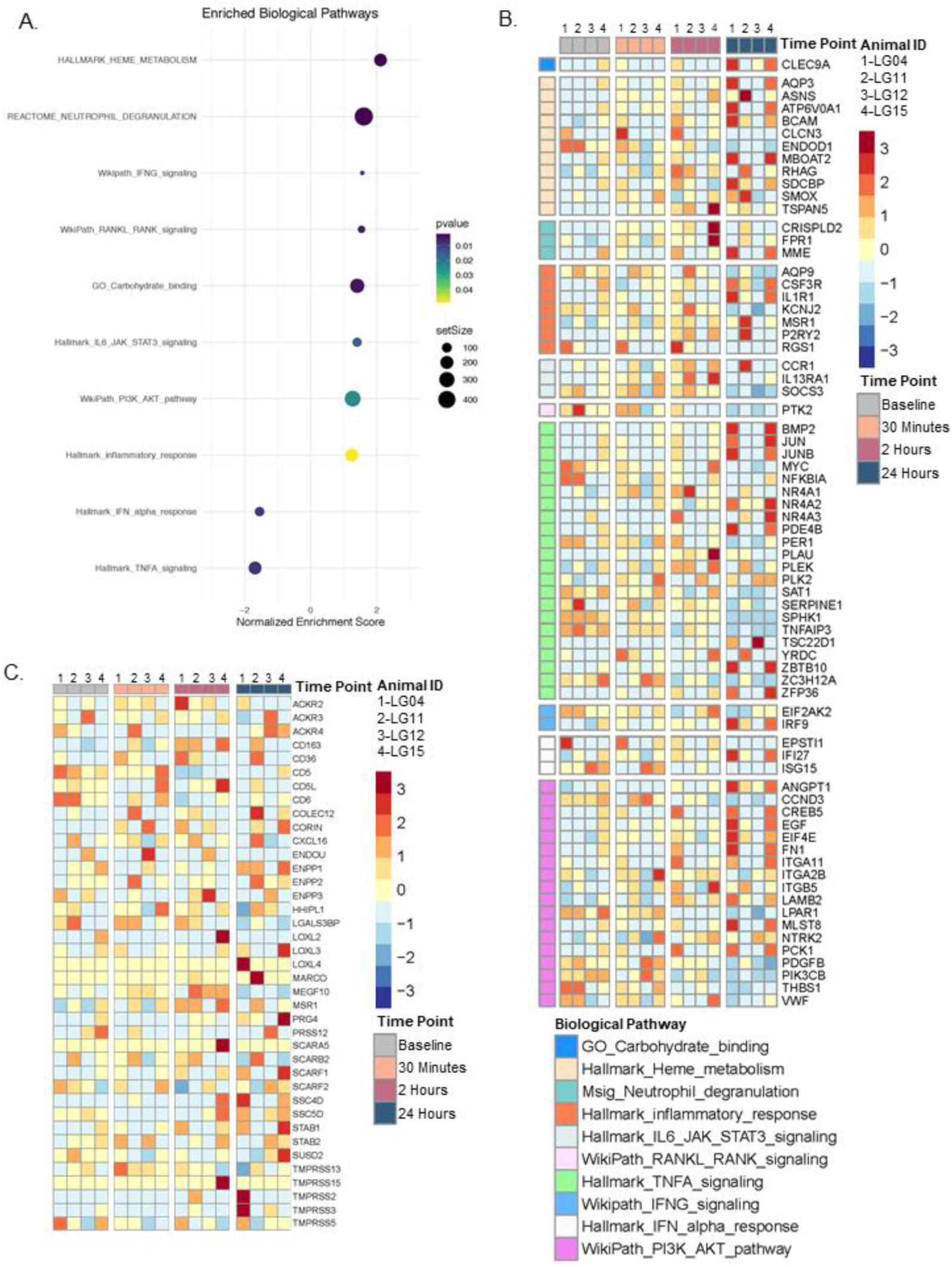
Biological pathways related to NK T cells are enriched upon anti-NKT antibody administration. (A). Normalized enrichment scores (NES) of relevant NKT-related pathways selected from an unbiased gene set enrichment analysis (GSEA). Color ramp represents NES significance defined by a Fisher’s Exact test. Dot size represents ‘setSize’, or number of genes contained within an *a priori*-defined biological pathway. (B) Composite heatmap of genes meeting a significance threshold (p < 0.05) at 30 minute-, 2 hour-, and 1 day-post NKT-antibody administration that are contained within significantly enriched pathways depicted in (A). Rows are scaled by z-score-normalized expression. (C) Heatmap of genes within the GO pathway “scavenger receptor activity”.

Targeted analysis of differentially expressed genes showed elevation of several lymphoid genes related to monocytes, granulocytes, B-cells, T-cells and NK-cells as early as 30 minutes after NKTT320 administration (Figure 6C). Notable among these were upregulation of the following genes: the low affinity inhibitory Fc gamma receptor FCGR2B involved in regulation of antibody production by B-cells and modulation of antibody-dependent effector function of myeloid cells (Nimmerjahn and Ravetch, 2008); Fc fragment of IgA receptor FCAR present on myeloid lineage cells and mediating phagocytosis and ADCC (Monteiro and Van De Winkel, 2003); C-type lectin-like receptor CLEC1B expressed on NK cells (Huysamen and Brown, 2009); CD14, CD163 expressed on monocytes and macrophages; IL-13 receptor alpha-1 chain (IL13RA1) involved in activation of JAK1, STAT3 and STAT6 induced by IL13 and IL4 (McCormick and Heller, 2015); the MCP-1 receptor CCR2 mediating monocyte chemotaxis (Deshmane et al., 2009); CD40 ligand (CD40L) expressed on iNKT and T-cells promoting DC-iNKT interactions and regulating B-cell function through CD4 help (Brennan et al., 2013); colony stimulating factor 3 (CSF3R) or G-CSF receptor controlling granulocyte maturation and function (Ward, 2007); and N-formyl peptide receptors (FPR1, FPR3) that are powerful neutrophil chemotactic factors (Dahlgren et al., 2020). Several inflammation-related and pattern-recognition receptor genes were elevated. These included IL1R1; extracellular and cytosolic receptors for bacterial flagellin, namely toll-like receptor 5 (TLR5), and neuronal apoptosis inhibitory protein NAIP that also acts as the sensor component of the NLRC4 inflammasome and promotes caspase-1 activation (Zhao and Shao, 2015); the carbohydrate sensing innate immune recognition ficolin FCN1 expressed in monocytes (Gout et al., 2010); and the C-type lectin receptor CLEC4E or MINCLE expressed on macrophages that recognizes fungal and mycobacterial ligands (Furukawa et al., 2013). Upregulation of the chemokine receptor genes CCR2 and CCR1 was concordant with elevated plasma levels of their respective ligands, CCL2 and MIP-1α, post NKTT320 administration (Figure 6 and Supplemental Table 1). Overall, upregulation of these genes was indicative of a broad stimulation of the innate immune system with facilitation of antigen presentation function.

Simultaneously an immune downmodulation effect of NKTT320 was evident by upregulation of several immune inhibitory and inflammation suppressive genes (Figure 6). These included the T-lymphocyte inhibitor PD-1 ligand 1 (CD274) (Freeman et al., 2000); the arginine metabolism enzyme ARG2 that can regulate inflammation and immunity (Asosingh et al., 2020, McGovern et al., 2017, Geiger et al., 2016, Dowling et al., 2021); the potent mitigator of inflammation, nucleotide-binding oligomerization domain protein NLRP12 that acts as a negative regulator of NF-κB and promotes degradation of NOD (Normand et al., 2018); the chemerin chemokine-like receptor CMKLR1 which binds the endogenous lipid mediator Resolvin E1 and actively regulates resolution of acute inflammation (Ohira et al., 2010), the suppressor of cytokine signaling SOCS3 involved in the negative regulation of cytokines such as IL6 that signal through the JAK/STAT pathway (Heinrich et al., 1998), and ROR1 that can inhibit the Wnt3a-mediated signaling pathway (Bainbridge et al., 2014). These data point to an inflammation suppressive effect of NKTT320-mediated iNKT activation acting in concert with activation of B-cells, monocytes, dendritic cells, and T-helper pathways. It is noteworthy that both activating and suppressive effects were detected in the first 24-hours of NKTT320 administration. The inhibitory signals may reflect a feed-back loop following initial activation.

Downregulation of genes such as PRF1, NCR1, GNLY, GZMB, GZMH, CD244 associated with T-lymphocyte and NK cytotoxicity (Figure 6) was unexpected due to the functional evidence for T-lymphocyte activation. Of these, CD244 or 2B4, which belongs to the signaling lymphocyte activation molecule (SLAM)-receptor family and is expressed on NK cells, as well as on some T-cells, monocytes and basophils, can serve as both an activating and inhibitory receptor (Eissmann et al., 2005).

We followed the targeted gene analysis with an unbiased analysis of differentially expressed genes post NKTT320 administration (Figure 7). Using Gene Set Enrichment Analysis against four pathway databases, an aggregate of 201, 332, and 423 enriched pathways were detected at 30 minutes, 2 hours and 24 hours respectively post NKTT320 administration (Supplemental Tables 3-4). Normalized enrichment scores of 10 biological pathways related to NKT-lymphocytes that were significantly altered on NKTT320 administration along with heatmaps of genes in these pathways reaching a significant threshold are shown (Figure 7A-B). Because of CD163 upregulation and upregulation of genes involved in phagocytosis we also evaluated genes in the scavenger receptor activity pathway (Figure 7C).

Several genes from the inflammatory response, heme metabolism, neutrophil degranulation, IL-6/JAK/STAT3, and PI3K/AKT pathways were increased within 24 hours of NKTT320 (Figure 7B). Among the upregulated genes, elevation of the CLEC9A gene in the carbohydrate binding pathway was noteworthy. Targeting antigens to the C-type lectin CLEC9A on DCs can induce strong humoral immunity and T follicular helper responses independent of adjuvant and is being explored as a vaccine strategy (Caminschi et al., 2008, Cueto et al., 2019). Upregulation of the heme metabolism pathway SDCBP gene encoding the syndecan binding protein or syntenin-1 is interesting as it regulates TGFβ1-mediated downstream activation (Hwangbo et al., 2016).

Downregulated genes in the pathway analysis included the cyclin family gene CCND3, platelet-derived growth factor PDGFB, PIK3CB, and LPAR1, a member of the G protein-coupled receptor superfamily with diverse biological functions including proliferation and chemotaxis. The functional effects of genes that were downregulated appeared to be both pro-activation as well as anti-inflammatory. Several of the downregulated genes in the TNFA signaling pathway were inhibitors of NF-κB activation and it is likely that their suppression would lead to increased activation. Examples include suppression of TNFAIP3 and ZC3H12A, an IL-1-inducible gene encoding the monocyte chemotactic protein-1-induced protein-1 (MCPIP1) that acts as a transcriptional activator but also suppresses NF-κB activation (Jura et al., 2012). Similarly, suppression of the sphingosine kinase 1 gene (SPHK1) has been shown to potentiate induction of RANTES (Adada et al., 2013). The clock gene PER1 regulates pro-inflammatory mediators, and its suppression can lead to increased CCL2 and IL6 (Sugimoto et al., 2014). On the other hand, downregulation of PIK3CB or PI-3 kinase subunit beta in the PI3K/AKT pathway can be instrumental in suppressing inflammation by prevention of AKT phosphorylation and inducing FOXO activation (Finlay and Cantrell, 2011).

### iNKT maintain proliferative ability and avoid anergy following NKTT320 treatment

One hurdle to therapeutic modalities of *in vivo* iNKT activation is the induction of iNKT anergy. Administration of soluble αGC *in vivo* results in rapid iNKT activation followed by subsequent iNKT anergy in response to further stimulation (Parekh et al., 2005). To test whether *in vivo* NKTT320 treatment induces anergy, we performed *in vitro* proliferation assays in PBMC from NKTT320-treated animals, measuring Ki67 and BrdU double-positive cells after a 6-day stimulation with αGC. iNKT of animals that had received NKTT320 did not display anergy. On the contrary they were more responsive to *in vitro* αGC stimulation and showed increased proliferation compared to a pre-NKTT320 time-point (Supplemental Figure 5A). Furthermore, iNKT-cells continued to expand in culture on αGC stimulation pre- and post NKTT320 treatment providing further evidence that the monoclonal antibody does not induce iNKT anergy (Supplemental Figure 5B). The increased *in vitro* iNKT expansion was specific to αGC stimulation and not seen with other stimuli (Supplemental Figure 5B). These results were reproducible at day 1 post NKTT320 in 3 of 5 animals assessed by proliferation assays (data not shown). We saw a comparable or greater increase in plasma IL1RA, IL-6 and MCP-1 after a second dose of NKTT320, providing further evidence for absence of anergy (Figure 8A). In one multiply dosed animal, iNKT proliferative ability temporally followed increases in NKTT320 plasma concentration after each dose (Figure 8B).

**Fig. 8.**
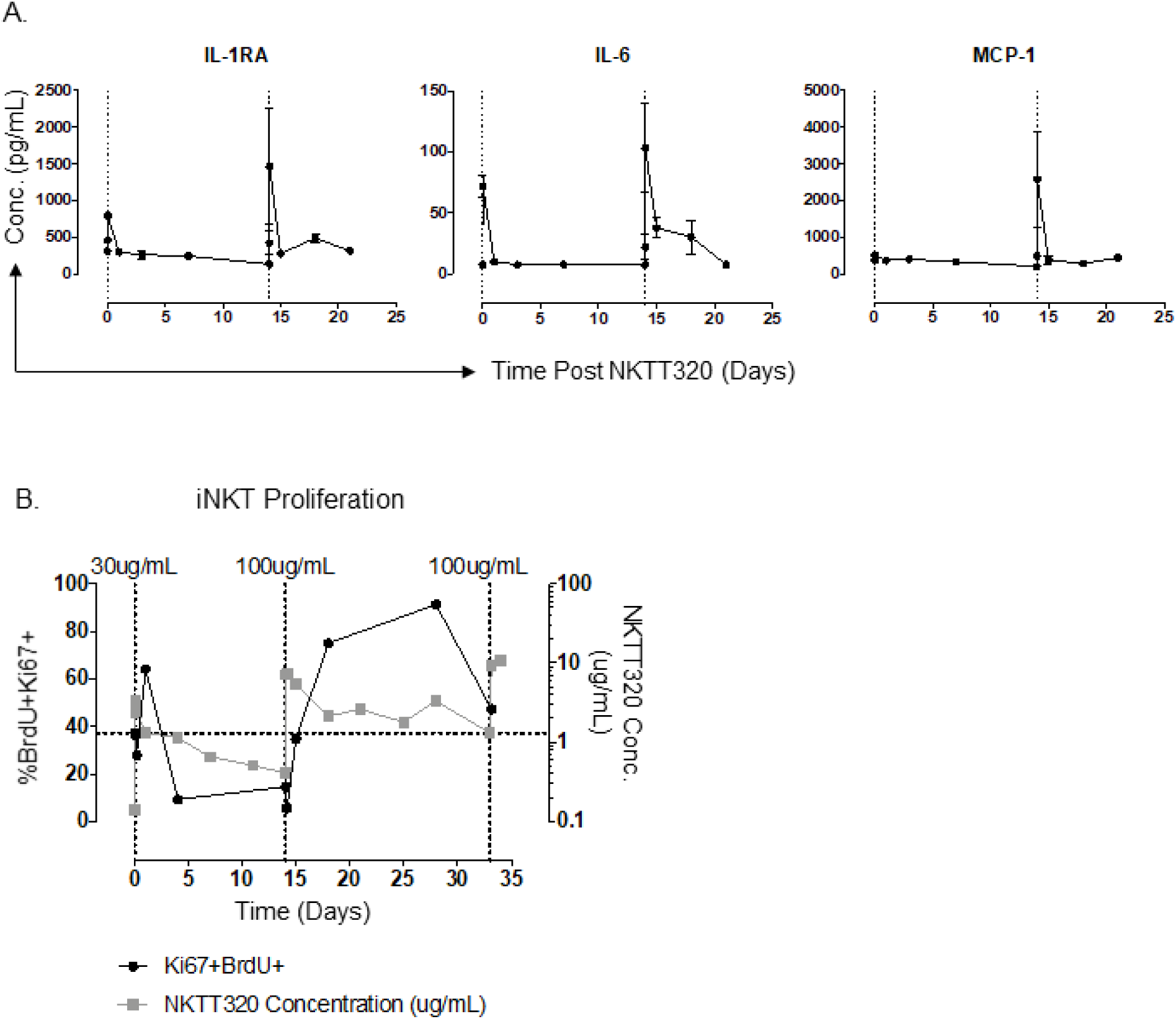
iNKTs retain proliferative capacity after NKTT320 indicating a lack of NKT anergy. (A) *Ex vivo* plasma luminex data from animals that received 2 doses of NKTT320 (n=4). (B) iNKT proliferation measured as BrdU+ Ki67+ double positives by flow cytometry. Proliferation data is compared to plasma NKTT320 concentration measured by NKTT320 ELISA. Vertical lines indicate the time and concentration of each dose. Horizontal line indicates baseline iNKT proliferation. Vertical lines indicate timing of dosing. Data in one multiply dosed MCM.

### Increased iNKT frequency in adipose tissue after NKTT320 administration

Our observations on the effect of NKTT320 were thus far confined to the examination of peripheral blood where we did not detect an increase in iNKT frequency. To investigate trafficking or effect on tissue iNKT, we examined iNKT frequency pre-and 14 days post-NKTT320 administration in lymph node (LN), bone marrow (BM), bronchoalveolar lavage (BAL), rectal mucosa (REC) and adipose tissue in a subset of animals. Adipose tissue is known to be a site of iNKT localization in humans (Lynch, 2014). Interestingly, we observed an increased frequency of adipose iNKT while iNKT frequency at other tissue sites was unchanged (Figure 9A-B). This suggests that either adipose-resident iNKT proliferate in response to NKTT320 treatment or iNKT are trafficking to adipose tissue following NKTT320 treatment.

**Fig. 9.**
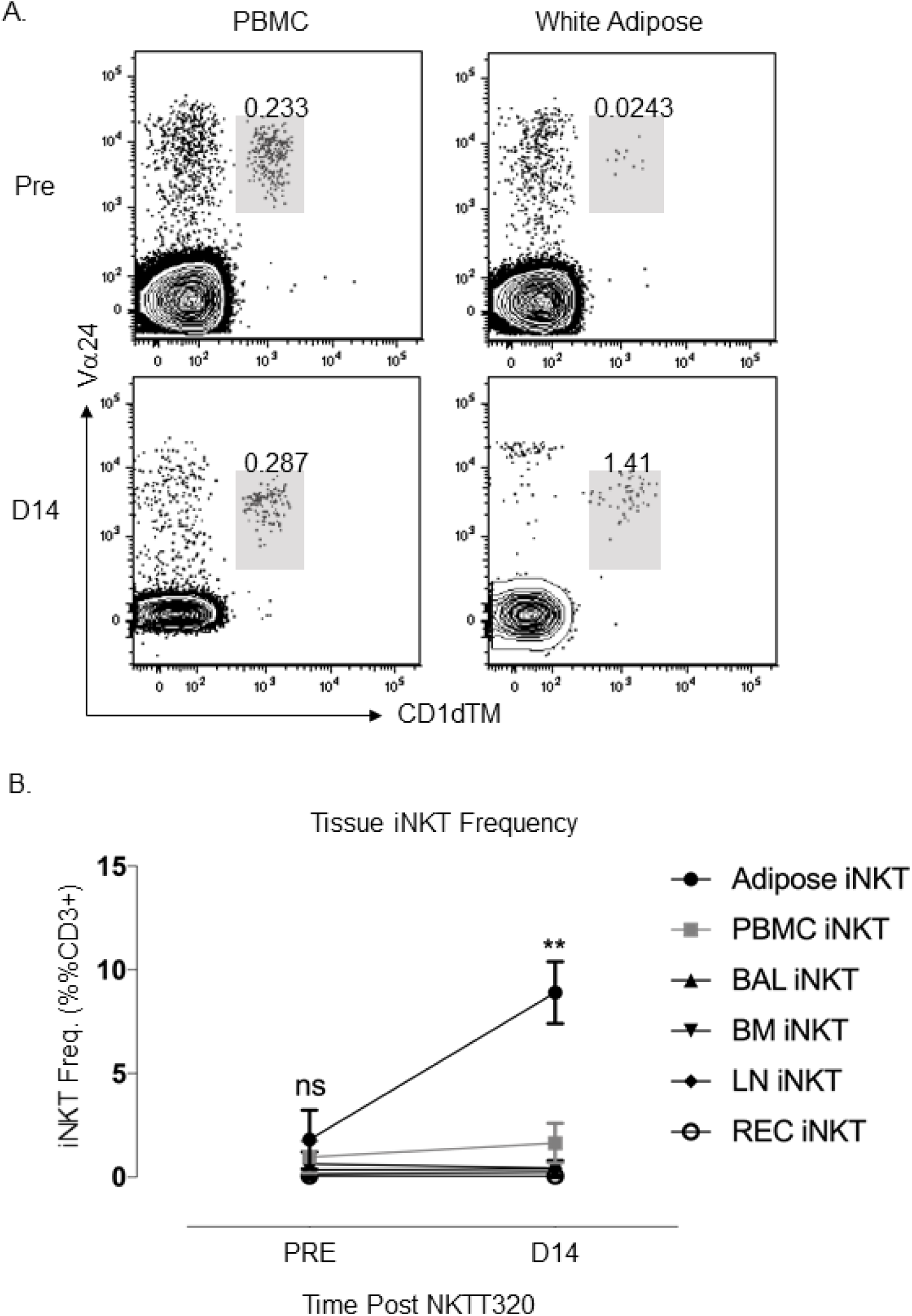
NKTT320 treatment *in vivo* results in trafficking of iNKTs to adipose tissue sites. (A) Representative plots of iNKT frequency pre and post NKTT320 treatment in white adipose and PBMC. (B) iNKT frequency in tissues and PBMC pre and 14 days post NKTT320 treatment (n=3). Paired t-test, **<0.01. ‘PBMC’-peripheral blood mononuclear cells; ‘BAL’-bronchoalveolar lavage; ‘BM’-bone marrow; ‘LN’-lymph node; ‘REC’-rectal mucosa.

## Discussion

This study is the first demonstration of sustained *in vivo* iNKT activation using the novel iNKT-activating humanized monoclonal antibody, NKTT320, that selectively binds with high affinity to the invariant NKT TCR in humans (Patel et al., 2020). We used the nonhuman primate model of MCM to extensively characterize the *in vitro* and *in vivo* effects of NKTT320 and assess its utility as an adjuvant and immunomodulatory tool. *In vitro*, NKTT320 showed dose-dependent iNKT-specific activation and increased cytokine production. *In vivo*, a single intravenous inoculation of NKTT320 was sufficient to rapidly induce iNKT activation that was sustained for up to 6 weeks without causing anergy. *In vivo* NKTT320-induced iNKT activation was associated with downstream activation of non-NKT immune cell subsets and iNKT trafficking to or proliferation within adipose tissue. Even though iNKT accounted for only 0.1-10% of circulating T-lymphocytes, iNKT activation had a profound amplification effect due to downstream effects on a wide range of immune cells. Our findings on *in vivo* iNKT activation and downstream effects on CD4^+^ T-lymphocytes, monocytes, dendritic cells and B cells, make NKTT320 a promising candidate for immunotherapy and vaccine adjuvant with translational potential.

*In vivo* NKTT320 administration led to iNKT-specific activation within 30 minutes of administration and was accompanied by increased iNKT polyfunctionality with Th1 and Th2 cytokine secretion, increased proliferative capacity and trafficking to adipose tissue. A greater stimulatory effect on CD4^+^ iNKT was observed as they were the major source of the cytokines contributing to increased polyfunctionality in mitogen-stimulated iNKT. An increase in IL-2 and IL-4 was followed by IFNγ and TNFα production. IL-2 and IL-4 are both known to regulate and promote T-cell differentiation into Th1 and Th2, respectively. Additionally, lymph node iNKT secreting IL-4 were recently shown to be a key mediator of humoral immunity, improving B-cell maturation and differentiation as well as antibody class switching (Gaya et al., 2018, Lee et al., 2015). Early increases in iNKT function were followed from day 3 onwards by increased cytokine secretion from non-iNKT T-lymphocytes, most notably IL-2, IL-17 and IL-22. Looking beyond the adjuvant potential of NKTT320, these Th17 cytokines are of particular interest in the context of HIV infection as they are known to play a major role in maintaining gut mucosal integrity.

NKTT320-induced activation had features that were distinct from glycolipid iNKT agonists. While NKTT320 activated iNKT within two hours of stimulation and reached peak response levels by 16 hours, αGC stimulation was slower to catch up requiring 48 hours to reach comparable levels. Additionally, NKTT320 resulted in a broader Th1 and Th2 response whereas αGC skewed towards Th1 alone. Whether this holds true *in vivo* remains to be seen. Few *in vivo* studies of iNKT activation using αGC in humans and macaques have described functional changes by ICS in response to mitogen stimulation. When described, increases in Th1 cytokine production, primarily IFNγ were reported (Fernandez et al., 2013). Although synthetic sphingolipid NKT agonists that drive Th2 responses are available (Bricard et al., 2010), NKTT320 may be advantageous because it fosters an environment of increased responsiveness directing polyfunctional responses to specific stimuli without the need for structural modifications. Furthermore, NKTT320 does not rely on CD1d-mediated antigen presentation for iNKT activation. This could be an advantage in disease settings such as tumors and HIV infection associated with CD1d downregulation. Another major advantage of NKTT320 is the absence of NKT anergy. Following *in vivo* treatment with αGC, iNKT rapidly become anergic and are unable to respond to further stimuli-either iNKT specific or general TCR stimulation (Parekh et al., 2005). The ideal tool for *in vivo* iNKT modulation would transiently activate iNKT while avoiding anergy. The fact that *in vivo* NKTT320 activates iNKT in the absence of anergy further underscores the promising nature of this antibody as an effective tool for *in vivo* immune manipulation.

Circulating iNKT frequencies were transiently reduced post NKTT320, which is consistent with previous studies of *in vivo* iNKT activation using αGC in pig-tailed macaques and humans (Fernandez et al., 2013, Giaccone et al., 2002, Woltman et al., 2009). It is important to consider that the decrease in circulating iNKT frequency could be the result of a masking effect due to NKTT320 binding to the same site as CD1dTM, or to activation and subsequent downregulation of the iNKT TCR rendering it difficult to detect iNKT. Another explanation is that iNKT trafficking to tissue effector sites after activation could also transiently decrease the frequency of circulating iNKT. Our observation of elevated iNKT frequencies in adipose tissue following NKTT320 treatment suggests this possibility. This is an interesting finding as adipose tissue has been recently recognized as an inflammatory site with significant involvement in metabolic syndromes (Saltiel and Olefsky, 2017). Additionally, adipose tissue is a known site of HIV/SIV reservoir (Damouche et al., 2015). The ability of NKTT320 to induce either proliferation or trafficking of iNKT to adipose tissue has implications for use in targeting the HIV/SIV reservoir. Further investigation of the tissue effects of NKTT320 are warranted.

Due to the unique regulatory function of iNKT-cells, iNKT activation influences downstream immune cells. Through multiple lines of evidence based on flow cytometry, Luminex and RNA-Seq data, we show profound and broad effects of NKTT320 on the innate and adaptive immune system, particularly on monocytes/macrophages. Increased levels of plasma IL-6, CCL2 and CXCL10 were detected as early as 30 minutes after NKTT320 antibody administration. A significant increase in circulating CD14^+^ monocyte frequency was observed at day 1 along with rapid upregulation of CD14, CD163 and CLEC4E gene expression within 30 minutes of antibody administration. Additionally, CCR2 (present on activated macrophages) gene expression was significantly upregulated at two hours in concert with significantly elevated plasma levels of its ligand CCL2. These data point to a rapid release of proinflammatory cytokines and chemokines along with mobilization of monocytes and activated macrophages initiated by NKTT320 administration. The downstream effects of NKTT320-mediated iNKT activation also included release of chemo-attractants involved in the recruitment of granulocytes, NK cells, and T-lymphocytes to sites of inflammation. Increase in chemokines involved in granulocyte recruitment included CCL11 (eotaxin) for eosinophil recruitment; CXCL8 (IL8) for neutrophil recruitment; CCL2 (MCP1) chemoattractant for basophils and monocytes; CXCL9-11 for chemoattraction of Th1 CXCR3-expressing T-cells to inflammatory sites; and CCL22 (MDC) chemoattractant for monocytes, DC, NK and T-lymphocytes (Bromley et al., 2008). Surprisingly, several genes in the IFNγ and TNFα signaling pathways and cytotoxic related genes were downregulated despite evidence of elevated plasma TNFα and IFNγ by Luminex. Although unexpected, the transcriptomics data represented a snap-shot of the first 24 hours whereas in the Luminex we saw plasma levels of TNF-α and IFN-γ peak only at 72 hours. mRNA and protein expression levels are often discordant for the same time-point during dynamic transitions, as was the case in the first 24 hours of NKTT320 administration which likely explains differences in Luminex and RNA-Seq data (Liu et al., 2016).

Several pattern recognition receptor genes expressed on dendritic cells and monocyte/macrophages were modulated by NKTT320. Notable among them was the C-lectin type receptor CLEC9a which is expressed on DCs and has been shown to efficiently induce humoral immunity and a T follicular helper response when antigen targeted to CLEC9a displays it to B-cells (Caminschi et al., 2008, Kato et al., 2015). Evidence of early DC activation was manifested by increased plasma IL-12 detection along with increased gene expression for a range of C-lectins associated with APCs. Additionally, gene expression of TLR5, a ubiquitous pathogen recognition marker found on DCs, and NAIP involved in bacterial flagellin recognition were upregulated. CD40 ligand (CD40L) gene expression was also upregulated likely on NKT and other CD4^+^ T-lymphocytes. A major pathway by which NKT promote DC/APC maturation is by upregulation of CD40L providing co-stimulation to DCs via CD40L and IFNγ, TNFα (Brennan et al., 2013). Moreover, NKT licensing of DCs for antigen cross-presentation can determine the type of the immune response. These data underscore the effect of NKTT320 on APCs which could lead to improved antigen presentation and potentiation of cellular immunity.

In addition to effects on APCs, we found that *in vivo* NKTT320 treatment had downstream effects on B-cells suggesting synergistic effects that likely improve humoral immunity. In the absence of a vaccine allowing a direct readout of humoral immune responses we found several indications of effects of NKTT320 treatment on B-cells. We found that B-cell proliferation was significantly increased, with Ki67 expression on circulating CD20^+^ B-cells reaching significantly higher levels at day 1 after antibody administration. Furthermore, increased B-cell proliferation paired with increases in IL-4 expression from iNKT within 2 hours of NKTT320 administration, suggests the potential for this antibody to improve humoral immunity. These findings were corroborated by the rapid upregulation of B-cell related genes FCAR, IGJ and FCGR2B. These data show the potential to harness the effect of iNKT using NKTT320 to improve APC and B-cell function ultimately improving antigen-specific antibody responses in the context of vaccination or infection.

Interestingly, our data show the potential of NKTT320 to modulate both inflammatory and anti-inflammatory responses. As illustrated in the schematic (Figure 10), side-by-side with activation, NKTT320 triggered several genes encoding markers of resolution of inflammation. Most prominent of the upregulated genes included the inflammation resolution genes NLRP12 and CMKLR1 (Fullerton and Gilroy, 2016, Ohira et al., 2010), the metabolic regulator ARG-2 (Dowling et al., 2021), and the inhibitory receptors FCGR2B and PD-L1 (Nimmerjahn and Ravetch, 2008, Freeman et al., 2000). In all, the concordant immune activation and anti-inflammatory responses induced by NKTT320 may mean that there is a potential *in vivo* for eliciting potent inflammatory responses without excessive or non-resolving immune activation. This remains to be tested in both long-term studies and in the context of vaccination or infection.

**Fig. 10.**
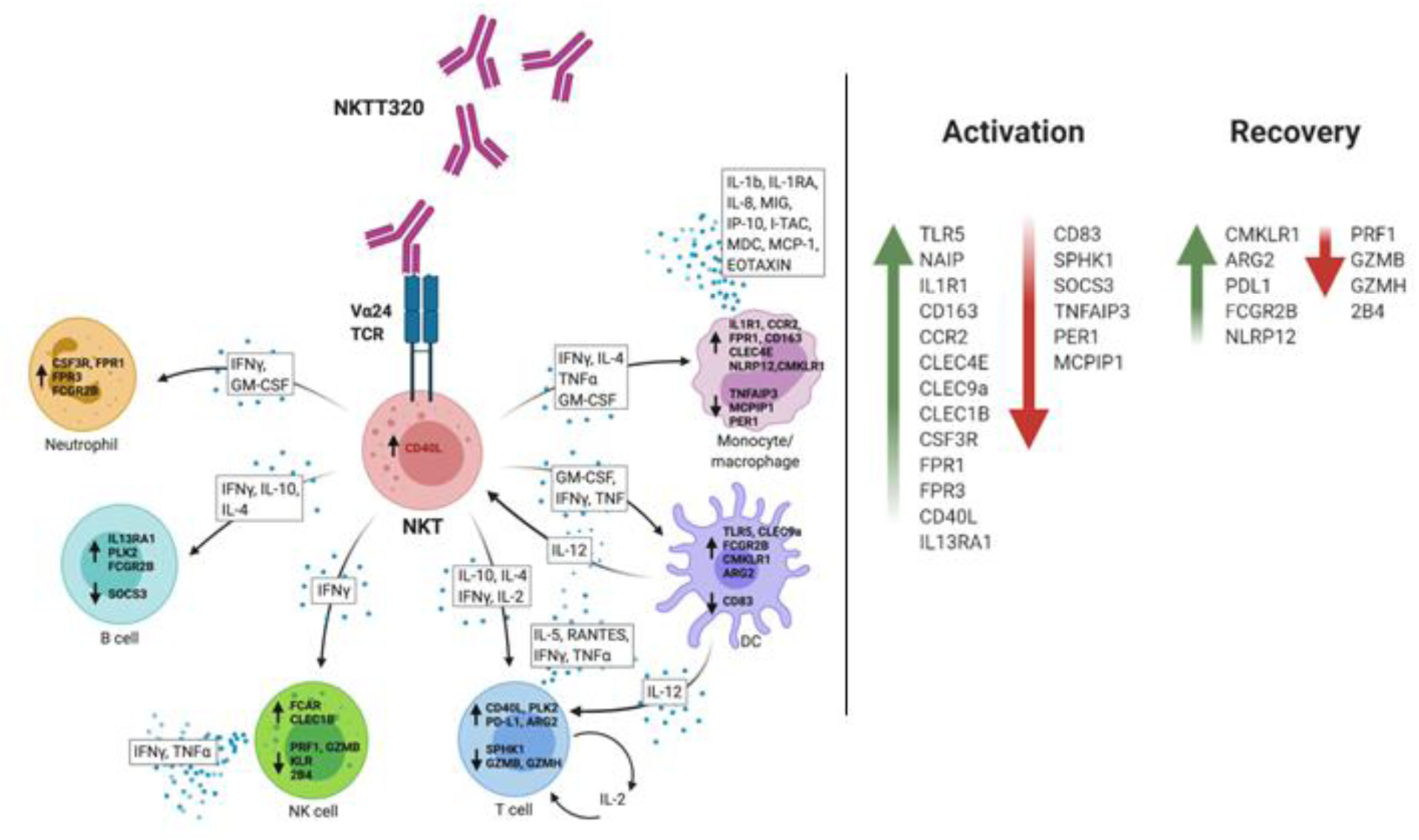
Schematic of NKTT320-mediated immune modulation. Schematic effect of NKTT320 on iNKT and downstream immune cell subsets as measured by Luminex, ICS, and RNA-Seq. Genes and secreted molecules are placed according to potential cellular sources.

In summary, the current studies detailed here investigate the use of NKTT320, an iNKT-specific humanized monoclonal antibody, for *in vivo* iNKT activation for the first time in the NHP model. We found, through multiple modes of investigation, that this antibody directly affects iNKT without inducing anergy and subsequently transactivates other immune subsets, most strikingly monocytes and macrophages. *In vivo* NKTT320 administration induces potent functional changes in both T-cell and non-T-cell subsets that may influence both innate and adaptive immunity. It also triggers a balanced immune response with induction of anti-inflammatory responses to balance immune activation. The long half-life of NKTT320 as well as detection of effects on iNKT functionality well beyond the duration of detectable plasma NKTT320 indicates that this therapeutic modality would be feasible in human studies or in a translational setting. NKTT320 is a promising iNKT activating agent with translational potential that should be studied further for efficacy as a vaccine adjuvant and immunotherapeutic tool.

## Materials and Methods

### Ethics statement for *in vivo* non-human primate studies

*In vivo* non-human primate studies were performed at New England Primate Research Center (NEPRC) (Boston, MA; 2013-2015) and Tulane National Primate Research Center (TNPRC) (Covington, LA; 2015-2018). This study was carried out in accordance with the Guide for the Care and Use of Laboratory Animals of the NIH. The protocol was reviewed and approved by the Institutional Animal Care and Use Committees (IACUCs) at NEPRC and TNPRC. The facilities also maintained an Animal Welfare Assurance statement with the National Institutes of Health, Office of Laboratory Animal Welfare.

### Animals and Study design

We longitudinally evaluated blood from adult male MCM (n=12) and tissues from a subset of these animals (n=3), pre and post NKTT320 treatment.

#### Single-dose pharmacokinetic studies

NKTT320 was administered by intravenous route at escalating doses in three groups: low (100μg/kg, n=6), mid (300μg/kg, n=3) and high (1000μg/kg, n=3). Samples were taken prior to NKTT320 administration, and 30 minutes, 2 hours, days 1, 3, 7, and weeks 2, 4, 6, 8, and 14 post treatment. Animals were released to the second phase of the study after the final time point.

#### Repeated dose studies

One animal received 3 doses of NKTT320—30μg/kg, 100μg/kg and 100μg/kg—at 2-week intervals.

### Processing of peripheral blood and tissues

Peripheral blood mononuclear cells (PBMCs) were isolated from MCM by density gradient centrifugation. In brief, peripheral blood was diluted 1:1 in PBS and layered over 90% ficoll at a 1:2 ratio. The gradient was then spun at 2,200 rpm for 45 minutes with the brake off. Cells were then isolated, washed and counted according to standard methods. PBMCs were used for phenotyping and functional assays. In a subset of experiments bone marrow (BM), bronchoalveolar lavage (BAL), peripheral lymph node (PLN), rectal mucosa, subcutaneous abdominal adipose tissue biopsies were taken. In brief, tissue lymphocytes were isolated as follows. BM was treated identically to peripheral blood, described above. BAL was strained, spun at 1,500 rpm for 5 minutes and counted. PLN lymphocytes were isolated using mechanical separation. Rectal mucosa lymphocytes were isolated by enzymatic digestion with EDTA followed by collagenase II. Remaining tissue was mechanically disrupted, strained and washed before counting. Similarly, adipose tissue was initially digested using collagenase II followed by mechanical disruption, straining out remaining tissue and washing the cells before counting. Tissue lymphocytes were used to assess iNKT frequency within tissues following activation with NKTT230.

### NKTT320 ELISA

NKT Therapeutics developed an enzyme-linked immunosorbent assay (ELISA) to determine NKTT320 concentration in monkey plasma or serum based on the detection of human IgG4. Briefly, 96 well plates were coated with monkey adsorbed goat anti-human IgG (1μg/mL) and incubated at 37°C for 2 hours or 4°C overnight. Plates were washed 3x with wash buffer (0.05% Tween 20 in 1X PBS). Wells were blocked using 3% BSA in 1X PBS for 2 hours at room temperature. Plates were washed as described above, diluted samples, standards and controls were added and incubated for either 2 hours at room temperature or overnight at 4°C. For antibody detection, 0.1μg/mL goat anti-human IgG Biotin (monkey adsorbed) was added to each well and incubated for 1 hour at room temperature. Plates were washed as described above, 100μL diluted streptavidin-HRP polymer (SPP) conjugate was added, and samples were incubated 1 hour at room temperature. Plates were washed as described above, 100μL of Tetramethylbenzidine (TMB) was added to each well, incubated in the dark for between 10-20 minutes and the reaction was stopped by adding 100μL stop solution to each well. Plates were read at 450nm within 30 minutes of adding stop solution on a Biotek synergy 2 (Winooski, VT) microplate reader and data was acquired and analyzed using Gen5 1.11.5 software, Excel and GraphPad/PRISM.

### Cells and reagents

#### CD1d transfectants

C1R cells (Rout et al., 2010) were used either loaded with αGC (Avanti Polar Lipids; Alabaster, AL) or alone to stimulate PBMC to compare a known iNKT specific stimulant with NKTT320 stimulation. C1R transfectanT-cells were used at a ratio of 1:10 C1R: PBMC for stimulation.

#### Alpha-galactosylceramide

αGC stock was received at 2mg/mL from Avanti Polar Lipids, Alabaster, AL and stored in 10μL aliquots in brown glass at −80°C. Aliquots of working solution of 20μg/mL in R10 medium (RPMI+Hepes+L-glutamine+10% fetal calf serum) were stored at −20°C until use. αGC was loaded onto C1Rd cells and used for iNKT stimulation at a final concentration of 100ng/mL.

#### NKTT320 monoclonal antibody

NKTT320 is a humanized monoclonal antibody developed by NKT Therapeutics (Sharon, MA) that binds to the CDR3 region of the Vα chain of the human iNKT TCR (Scheuplein et al., 2013, Patel et al., 2020, Truneh, 2013). NKTT320 is the sister antibody to NKTT120 (Field et al., 2017, Scheuplein et al., 2013). Briefly CD1d knock out mice, which lack iNKT-cells, were immunized with cyclic peptides from the CDR3 loop of the TCRα and screened for antibodies. One mouse monoclonal antibody designated 6B11, was identified as specific for human and non-human primate but not rodent iNKT-cells. 6B11 has demonstrated the ability to identify, purify, activate and expand iNKT-cells (Exley et al., 2008). To permit the clinical evaluation the murine mAb, 6B11, was humanized and de-immunized using the “Composite Human Antibody™ Technology” (Antitope Ltd., Cambridge, UK) to a stabilized IgG4 activating antibody NKTT320. NKTT320 was evaluated by Surface Plasmon Resonance (SPR) assays to determine the binding affinity to soluble human iTCR. NKTT320 binds specifically and selectively to the human iTCR with a K_D_ of approximately 44 nM. Measurement of binding affinity and functional characterization of NKTT320 was performed using recombinant human invariant TCR and cells. NKTT320 was produced by Antitope LTD (Cambridge, UK) in a transfected CHO-M cell line provided by Selexis sa (Chemin des Aulx Switzerland).

#### Monoclonal antibodies for flow cytometry

Monoclonal antibodies were used for phenotyping and flow cytometric functional assays. Antibody clone, vendor and panel information can be found in Supplementary Table 5.

### iNKT Detection

CD1dTMs loaded with αGC conjugated to either APC or BV421 were obtained from the NIH Tetramer core. These loaded tetramers were titrated and used at the optimal titrated volume. iNKTs are identified based on co-expression of CD1d-TM and Vα24 as described previously (Rout et al., 2010). Briefly, isolated cells were washed, incubated with titrated volumes of CD1dTM at room temperature for 30 minutes followed by addition of the remaining antibodies in the staining cocktail for 20 minutes. Standard flow cytometric protocols were used for the remaining surface or intracellular cytokine staining panels. A minimum of 2 million PBMCs were used for flow cytometric staining to allow for at least 200,000 lymphocyte events to be captured to visualize rare iNKT populations. All flow cytometry panels were run on either BD LSR-II or BD Fortessa instruments (BD, Franklin Lakes, NJ) by the TNPRC Flow Cytometry Core.

### *In vitro* iNKT stimulation

*In vitro stimulation with soluble NKTT320:* Peripheral blood mononuclear cells (PBMC) isolated from 3 MCM were incubated with escalating concentrations (0.1, 1, 10 and 25ug/mL) of NKTT320 diluted in R10 media (RPMI/1%Hepes/10%FBS). Cells were incubated overnight at 37°C and harvested for Flow Cytometry staining.

#### In vitro stimulation with cross-linked NKTT320

PBMC were stimulated with 200ng/mL NKTT320 cross-linked with goat-anti-mouse (GAM) IgG Fab2 fragments. Briefly wells were first coated with GAM-IgG at 2.5ng/mL (SeraCare, formerly KPL, Gaithersburg, MD) in 50mM TRIS buffer (pH=8.6), 1mL per well, and incubated for 1 hour at 37°C. Wells were then washed 3x with PBS and 1mL/well 200ng/mL NKTT320 in PBS was added. Plates were then incubated another hour at 37°C, washed and 1 million PBMC in R10 media were added to each well. Cells were then incubated 48 hours, harvested for flow cytometry; supernatants were harvested for Luminex assay.

#### In vitro stimulation with αGC

PBMC were stimulated with 100ng/mL αGC loaded onto C1Rd cells. C1Rd cells were irradiated at 10,000 rads (100 Gy) and mixed with PBMCs at a ratio of 1:1000. Finally, 100ng/mL αGC was added and plates were incubated 48 hours. Cells were then harvested for flow cytometry and supernatants were harvested for Luminex assay.

### Proliferation Assays

*In vitro* proliferation assays were conducted to assess iNKT proliferative ability and anergy following multiple *in vivo* doses of NKTT320. PBMC from days 0, 1, 7, 14 following each dose of NKTT320 were assessed. One million PBMC per condition were stimulated with either R10 media, aGC or staphylococcal enterotoxin B (SEB) (Sigma, Saint Louis, MO). Cells were seeded at 0.2M per well in a 96-well plate and brought up to 200μL/well with R10, 100ng/mL αGC or 200ng/mL SEB. Plates were incubated at 37°C for 5-6 days and 10uM Bromo-2’-deoxyuridine (BrdU, vendor) was added to each well 24 hours prior to staining. On the 6-7^th^ day of incubation, cells were pooled by time-point and stimulation condition and analyzed by flow cytometry.

### Luminex

Luminex technology was used to assess plasma chemokine and cytokine levels following NKTT320 treatment in 12 MCM. We used the Invitrogen (Carlsbad, CA) magnetic bead Monkey Cytokine Magnetic 29-Plex Panel covering the following analytes: EGF, HGF, VEGF, MIG, RANTES, Eotaxin, IL-8, GM-CSF, TNF alpha, IL-1 beta, IL-2, IL-4, IL-5, IL-6, IL-10, IL-12, MIP-1 alpha, IL-17, MIP-1 beta, IP-10, IL-15, MCP-1, G-CSF, IFN gamma, FGF-Basic, IL-1RA, MDC, MIF, I-TAC. Final reactions were read on a Bio-Plex® 200 System (Bio-Rad Laboratories, Hercules, CA) results were calculated using Bio-Plex Manager™ Software v6.2 (Bio-Rad) by the TNPRC Pathogen Quantification and Detection Core.

### RNA-Seq

#### Sample preparation for RNA-Seq

RNA was extracted from snap frozen, unfractionated PBMCs using the Qiagen RNeasy Plus Mini Kit (Qiagen 74134). Briefly, cells were first thawed and pelleted, then lysed following the kit protocol. gDNA was removed using a gDNA Eliminator spin column, and remaining flow through was added to a RNeasy spin column to bind RNA. The spin column/RNA was washed, RNA was eluted and stored at −80°C. Later, samples were concentrated using Zymo Clean & Concentrate-5 Kit (R1013). Briefly, RNA was thawed, bound, and added to the Zymo-Spin IC Column. Then RNA bound to the column was washed, eluted, aliquoted and stored at −80°C. Sample concentration was analyzed on the Qubit Fluorometer using the Qubit RNA BR Assay Kit. Samples were analyzed on the Tape Station at the Sequencing Core at CTRII Tulane University to determine RNA integrity number (RIN). Samples were then submitted for RNA-Seq analysis using a high output, and single read 75 cycle run.

#### RNA-Seq data analysis

RNA-seq for Eukaryotes Analysis v3 was used for sequencing analysis and built by the Banana Slug Genomics Center at the University of California Santa Cruz. First, an Illumina sequencer at the Tulane School of Medicine Genomics Core was used to construct raw sequencing reads which were checked for quality and contaminants using FastQC. Next, Trimmomatic was used to trim adapter sequences and primers from the sequencing reads (Bolger et al., 2014). Removal of polyA tail, polyN, and read segments with a quality score below 28 was accomplished by using PRINSEQ (Schmieder and Edwards, 2011). Following trimming, any reads of length less than 20bp were removed. Quality check was repeated, and high-quality reads were then mapped to the GRCh37/hg19 reference genome using STAR (Kim et al., 2013, Dobin et al., 2013) with NCBI RefSeq (O’Leary et al., 2016) annotated genes transcriptome index data. Raw read counts were normalized across all samples and then used for differential expression analysis using DESeq (Anders and Huber, 2010) and edgeR (Robinson et al., 2010) separately. Genes related to the immune system and with p-values of <0.05 from the edgeR pipeline were identified and categorized by major cell type, which were then plotted on individual heatmaps. Heatmaps were constructed using log_2_ fold change data and using the ‘pheatmap’ package in R (Kolde, 2019).

#### Gene Set Enrichment Analysis

GSEA was performed with the R package ‘ClusterProfiler’ at default parameters. Enrichment scores were calculated against the following 4 pathway databases containing a priori-defined gene sets: Gene Ontology (GO) database, Hallmark (H) and Curated (C2) gene sets of the Molecular Signature database (MsigDB), and WikiPathways. Gene sets significantly enriched in the datasets (p < 0.05) were subsequently curated for those relevant to NKT cell biological function. Enrichment plot was generated with the R software package ‘ggplot2’ and heatmaps with the ‘pheatmap’ package.

### Data Analysis

All flow cytometry data were analyzed using FlowJo version 9.9 (Ashland, Oregon). Cytokine polyfunctionality analyses were done using simplified presentation of incredibly complex evaluations (SPICE) (Roederer et al., 2011). Graphs were generated, and t-tests were performed using GraphPad/PRISM (LaJolla, CA). Statistical analyses were performed in GraphPad/PRISM using the non-parametric Wilcoxon Signed Rank test or paired t-test.

## Supplemental Materials

Supplemental materials include five supplemental figures referenced in text, three supplemental tables covering: additional in vivo Luminex data, significantly enriched pathways for the GSEA, and flow cytometry antibody clone/fluorochrome information. Finally, we submit two excel files containing data associated with the RNA-Seq analysis.

## Author Contributions

Designing research studies: AK, NB; conducting experiments: NB, SY, NR, DT, ES, TFS, LS; acquiring data NB, NR, DT, SY, ES, TFS, LS; analyzing data: NB, MF, JM, RS, AK; providing reagents: RS; and writing the manuscript: NB, AK.

## Acknowledgments

We would like to acknowledge TNPRC Veterinary Medicine Staff for caring for the animals, TNPRC Flow Cytometry Core for acquiring flow cytometry data, and the TNPRC Pathogen Detection and Quantification Core for reading and analyzing the Luminex plates. We thank Cathy Flemington and Alanna Wanek in the Tulane Center for Translational Research in Infection & Inflammation NextGen Sequencing Core for technical assistance for the RNA-Seq studies. We would like to acknowledge Dr. Mark Exley for thoughtful discussion. Study was funded by NIH/NIAID R01 AI102693 (AK), R21 AI145642 (AK) and P51 OD011104. The authors would like to thank the NIH Tetramer Core facility for provision of the CD1dTM conjugated to both APC and BV421.

