## Supplemental information for "Adjuvant and immunomodulatory potential of *in vivo* Natural Killer T (NKT) activation by NKTT320"

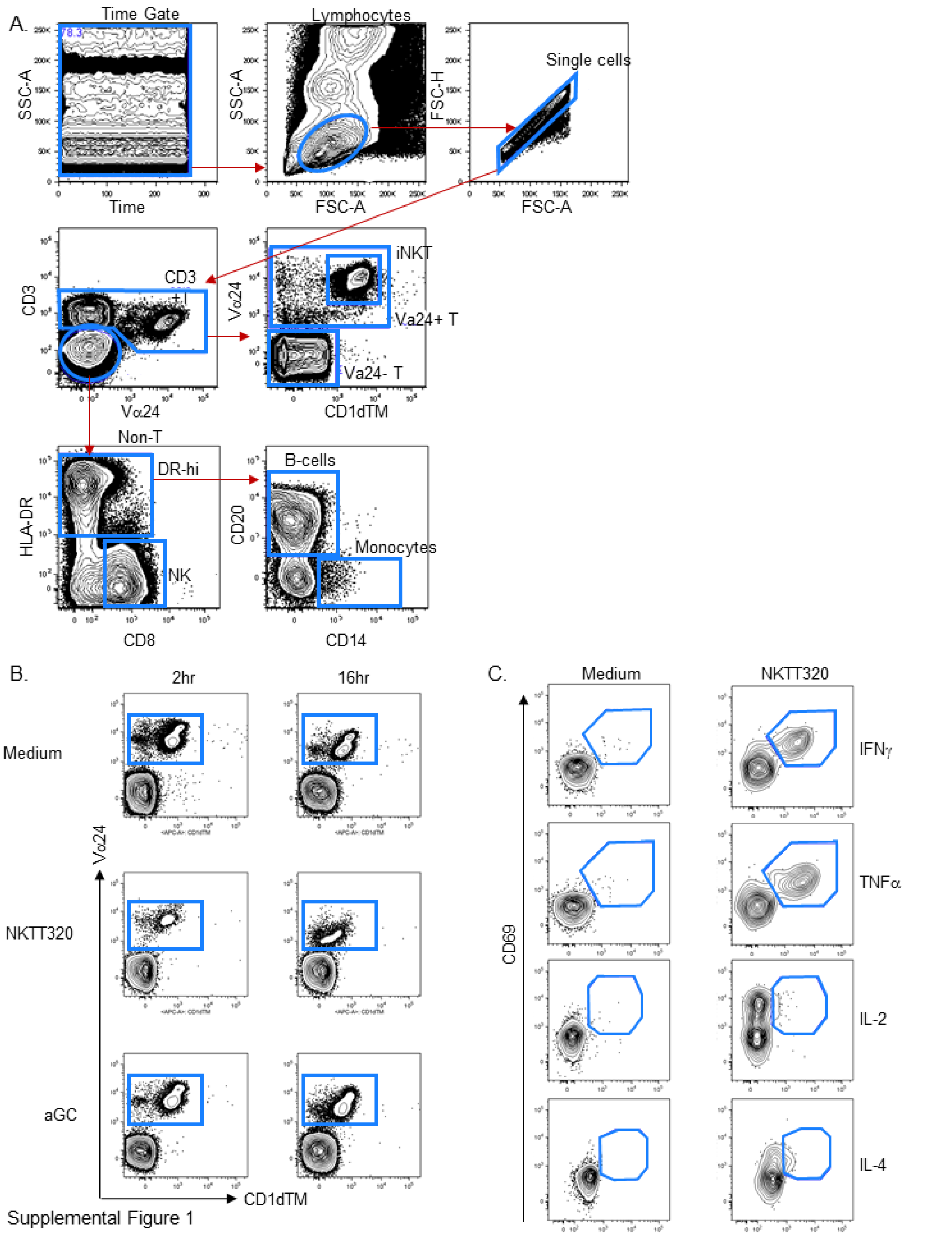


Supplemental Figure 1

**Supplemental Figure 1. Overall gating strategy.**
(A)General gating strategy for flow cytometry analysis showing immune subsets analyzed. (B) Representative plots of iNKT populations analyzed for cytokine expression comparing αGC and NKTT320 in Figure 2. (C) Representative plots of cytokine staining in Figure 2.

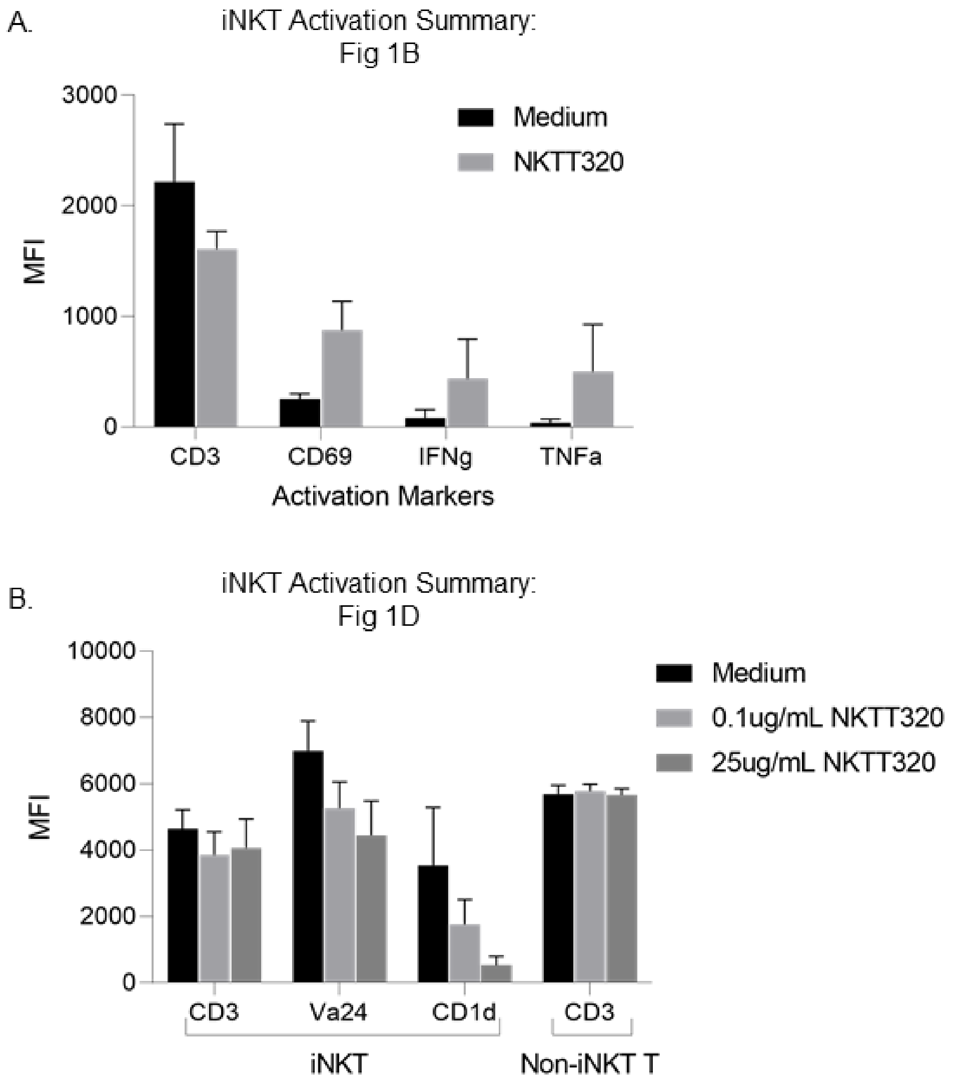


**Supplemental Figure 2. Summary data of iNKT activation markers *in vitro*.**
(A) Summary data of *in vitro* iNKT activation presented in Figure 1B (n=2). (B) Summary data of *in vitro* iNKT activation presented in Figure 1D (n=3).

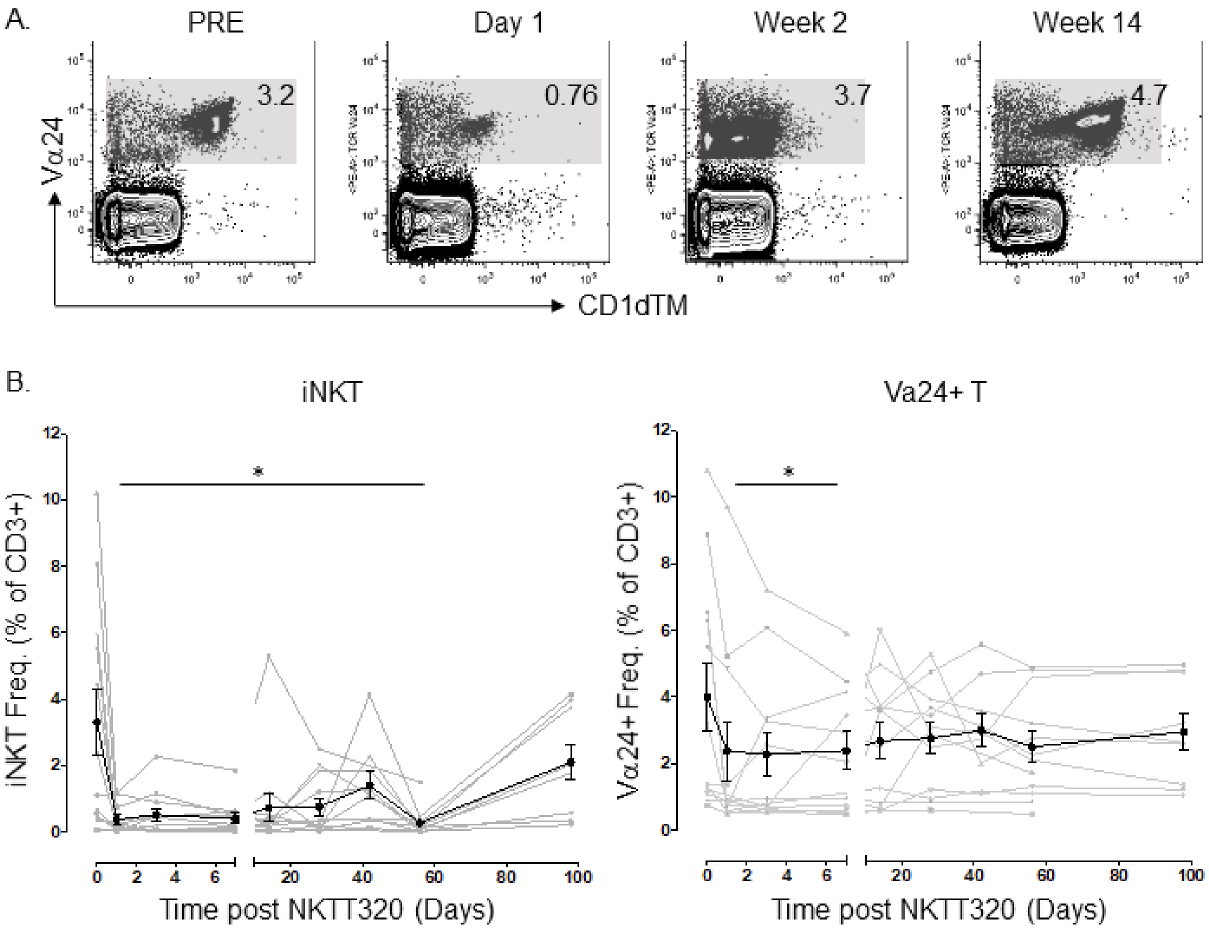


**Supplemental Figure 3. iNKT frequency following in vivo NKTT320 administration.** (A) Representative plots of total Va24+ frequency are shown with the population of interest shaded in gray. (B) iNKT and total Va24+ T-cell frequency variation measured by flow cytometry following NKTT320 treatment is shown in 11 MCM. Mean frequency is shown in black and individual animals are shown in gray. Error bars denote SEM.

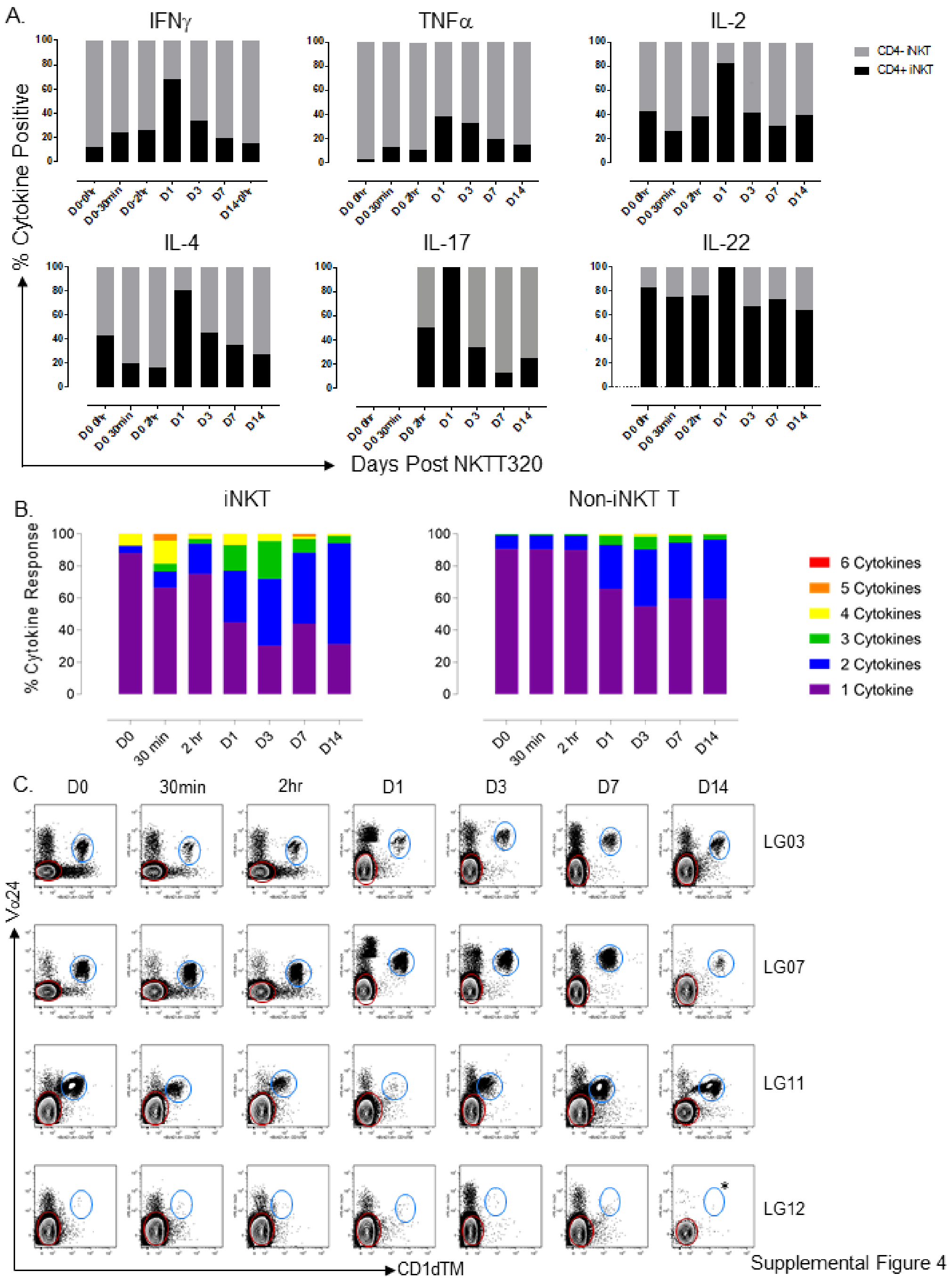


**Supplemental Fig. 4. NKTT320 rapidly induces functional changes in T-cell subsets.**

(A) Relative cytokine contribution from CD4+ versus CD4– iNKTs stimulated with PMA/ionomycin (n=7). Stacked columns showing proportion of cytokine secreted by CD4+ iNKT and CD4– iNKT at each time-point post NKTT320 administration. (B) Proportion of iNKT vs Non-iNKT T-cells responding with one or more cytokines after stimulation with PMA/ionomycin (n=4). (C) Plots of iNKT (blue) and Va24- (red) indicating the two populations analyzed for cytokine expression in figure 4. iNKT populations were only analyzed if there were greater than 10 events and a discrete population was visible. * indicates exclusion due to insufficient events.

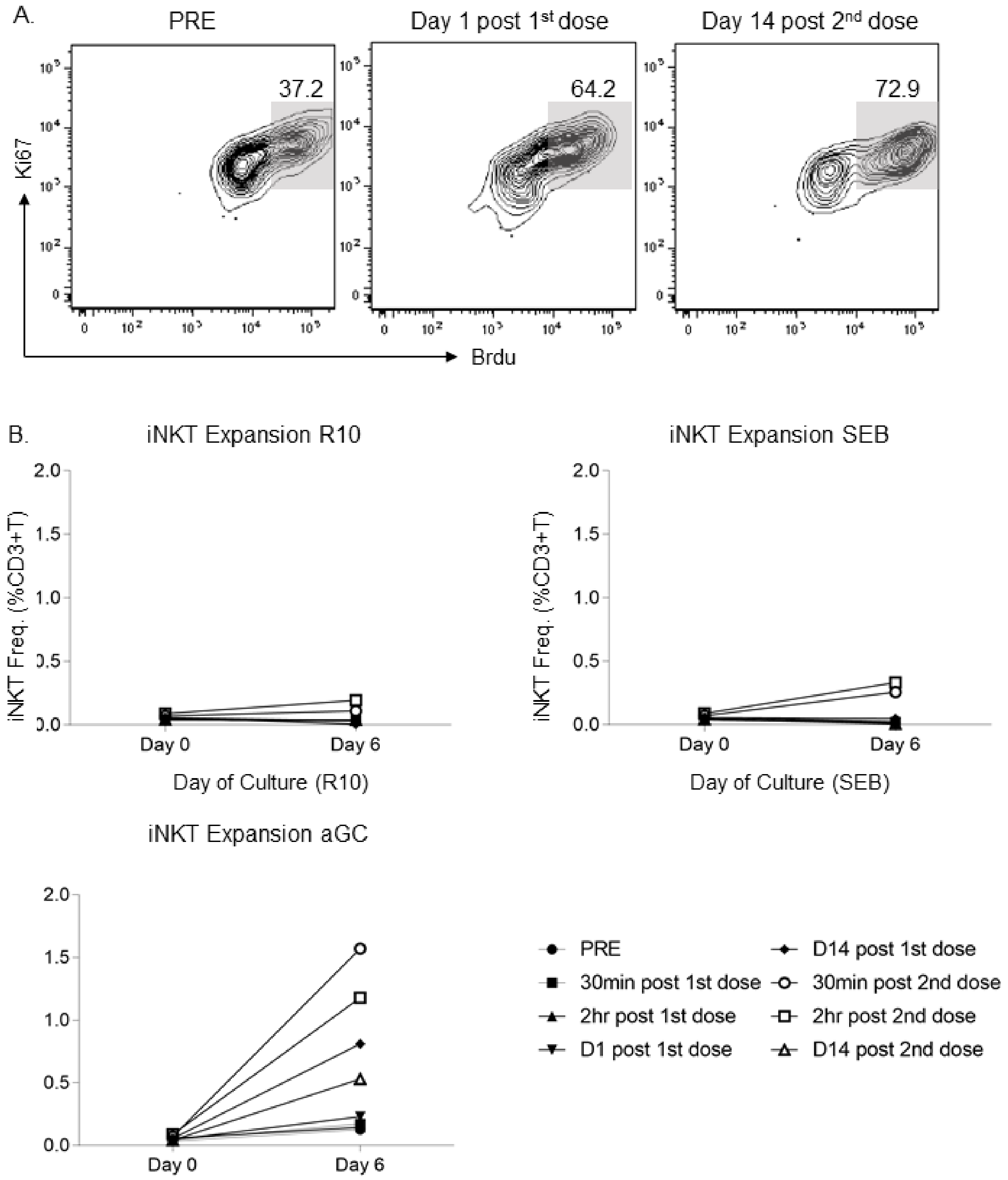


**Supplemental Fig. 5. iNKT frequency following 6-day culture with media, SEB, or αGC.**

(A) Representative plots of iNKT proliferative capacity after stimulation with aGC, pre-NKTT320, D1 post 1st dose and D14 post 2nd dose. Data from *in vitro* proliferation assays performed in 1 multiply dosed MCM. (B) iNKT expansion after 6 day culture with R10 media, SEB, and αGC at the following time points: pre-NKTT320, 30min, 2hr, D1 and D14 post 1st dose; and 30min, 2hr, and D14 post 2nd dose. Data from *in vitro* proliferation assays performed in one multiply dosed MCM.

Supplemental Table 1. In vivo Luminex summary in plasma post NKTT320 treatment.

| Analyte | Plasma mean (range) concentration in pg/mL (n=12) | | | Responding Animals | |
|---|---|---|---|---|---|
| | Baseline | Peak | p[1] | Time to peak (Hours) | Responders (n)[2] |
| IL-5 | 0.26 (0.09-0.38) | 0.54 (0.16-1.01) | **0.008** | 0.5 | 6 |
| IL-17 | 5.65 (1.37-9.61) | 6.53 (2.35-15.78) | 0.125 | 0.5 | 4 |
| G-CSF | 101.63 (27.40-203.76) | 526.96 (176.10-1979.41) | 0.250 | 0.5 | 5 |
| CCL3 (MIP-1$\alpha$) | 22.8 (9.44-34.75) | 46.78 (24.11-138.00) | 0.125 | 0.5-2 | 6 |
| CCL22 (MDC) | 616.41 (191.40-1153.25) | 1085.70 (748.40-1481.79) | **0.004** | 0.5-2 | 7 |
| VEGF | 0.199 (0.03-0.53) | 0.91 (0.16-3.62) | 0.063 | 2 | 8 |
| CXCL10 (IP-10) | 4.24 (0.48-10.54) | 8.52 (2.21-17.66) | **0.008** | 2 | 9 |
| IL-6 | 1.44 (0.15-3.17) | 25.11 (1.92-80.41) | **0.008** | 2 | 12 |
| CCL4 (MIP-1$\beta$) | 24.042 (8.45-47.78) | 35.97 (3.00-118.20) | **0.031** | 2 | 8 |
| IL-15 | 11.84 (2.71-26.31) | 83.72 (7.86-359.90) | 0.125 | 2 | 5 |
| CXCL11 (I-TAC) | 309.29 (148.04-631.81) | 349.38 (239.18-499.66) | **0.030** | 2 | 10 |
| IL-1RA | 244.98 (29.62-734.87) | 408.09 (116.74-885.19) | **0.017** | 2 | 9 |
| CCL11 (Eotaxin) | 350.06 (181.51-650.31) | 416.98 (211.71-611.00) | **0.001** | 2 | 10 |
| CCL2 (MCP-1) | 370.81 (148.34-771.36) | 590.38 (320.60-1050.15) | **0.0004** | 2 | 11 |
| IL-12 | 819.50 (353.16-1972.34) | 1011.23 (561.72-1358.68) | **0.0001** | 2 | 12 |
| IL-10 | 0.71 (0.14-1.26) | 1.62 (0.69-2.45) | **0.031** | 2-24 | 6 |
| CXCL8 (IL-8) | 11.18 (3.49-25.86) | 21.52 (9.70-41.27) | **0.047** | 2-24 | 8 |
| IL-4 | 4.026 (2.14-5.93) | 7.54 (2.14-12.64) | 0.125 | 2-72 | 8 |
| IL-1$\beta$ | 1.77 (0.09-2.99) | 5.64 (1.02-20.49) | **0.023** | 24 | 5 |
| EGF | 12.65 (0.48-50.24) | 13.23 (1.98-28.15) | 0.266 | 24 | 8 |
| FGF-basic | 13.20 (0.46-40.95) | 19.14 (2.79-59.08) | **0.001** | 24 | 9 |
| CXCL9 (MIG) | 40.16 (4.45-118.47) | 44.88 (16.26-118.47) | **0.031** | 24 | 5 |
| HGF | 64.20 (8.22-135.99) | 100.36 (27.17-186.18) | **0.004** | 24 | 7 |
| IL-2 | 131.89 (34.76-288.79) | 265.45 (86.57-1116.24) | **0.010** | 24 | 12 |
| MIF | 307.953 (37.69-1892.66) | 338.54 (103.54-1119.92) | 0.077 | 24 | 11 |
| CCL5 (RANTES) | 2531.00 (697.77-6678.73) | 5870.65 (1260.36-13983.48) | **0.007** | 24 | 9 |
| GM-CSF | 1.51 (1.32-1.72) | 2.3 (1.53-4.76) | **0.008** | 24-72 | 6 |
| TNF-$\alpha$ | 7.17 (2.66-9.98) | 12.46 (4.21-19.37) | **0.023** | 72 | 7 |
| IFN-$\gamma$ | 10.22 (0.65-17.18) | 28.43 (17.18-90.03) | **0.004** | 72 | 9 |

[1]Wilcoxon signed-rank test; p-value <0.05 shown in bold.
[2]Responding animals were defined as having ≥1.2-fold increase from baseline to peak.

Supplemental Table 2. RNA-Seq data associated with Figure 8. See Supplemental Data.

Supplemental Table 3. Significantly enriched pathways in four gene databases.

| Database | Number of Pathways | | |
|---|---|---|---|
| | 30 minute | 2 hour | 24 hour |
| Msig Hallmark | 3 | 3 | 4 |
| Msig Curated | 167 | 250 | 323 |
| WikiPathways | 15 | 47 | 59 |
| Gene Ontology | 16 | 32 | 37 |

Supplemental Table 4. RNA-Seq data associated with Figure 9. See Supplemental Data.

Supplemental Table 5. Antibody and clone information.

| Antibody | Fluorochrome | Clone | Vendor |
|---|---|---|---|
| Brdu | FITC | B44 | BD |
| CD11c | Alexa 488 | 3.9 | BioLegend |
| CD123 | PE-Cy7 | 7G3 | BD |
| CD14 | Alexa 700 | TuK4 | Caltag/Invitrogen |
| CD14 | PE-CF594 | M5E2 | BioLegend |
| CD1dTM | APC | aGC Loaded | NIH Tetramer Core |
| CD1dTM | BV421 | aGC Loaded | NIH Tetramer Core |
| CD20 | APC-Cy7 | L27 | B/D |
| CD20 | BV711 | 2H7 | BD |
| CD20 | BV605 | 2H7 | BD |
| CD3 | V500 | SP34-2 | Pharmingen |
| CD3 | APC-CY7 | SP34-2 | BD |
| CD4 | QDot 655 | T4/19Thy5D7 | custom/NHP Resource |
| CD4 | BV650 | OKT4 | BioLegend |
| CD45 | BV605 | D058-1283 | BD |
| CD45 | PE-CY7 | D058-1283 | BD |
| CD45 | V450 | D058-1283 | BD |
| CD69 | ECD | TP1.55.3 | Beckman/Coulter |
| CD69 | PE-CF594 | FN50 | BD |
| CD8 | Qdot 605 | T8/7Pt-3F9 | custom/NHP Resource |
| CD8 | AL 700 | RPA-T8 | BD |
| CD95 | BV421 | DX2 | Pharmingen |
| HLA-DR | PE-Cy7 | L243(G46-6) | BD |
| HLA-DR | PacBlue | L243 | biolegend |
| IFN-γ | BV711 | B27 | BD |
| IL-17 | PCP-Cy5.5 | eBio64DEC17 | eBioscience |
| IL-2 | BV605 | MQ1-17H12 | BioLegend |
| IL-22 | APC | IL22JOP | ebioscience |
| IL-4 | AL488 | 8D4-8 | biolegend |
| Ki67 | PerCP-Cy5.5 | B56 | BD |
| LIVE/DEAD | 405nm excitation | NA | Invitrogen |
| TNFα | AL 700 | MAb11 | BD |
| Va24 | PE | C15 | Beckman/Coulter |
